# Peritoneal cavity-derived GATA6^+^ macrophages inhibit fibrosis through IL33 in endometrium

**DOI:** 10.1101/2025.02.02.636170

**Authors:** Lijie Yin, Jingman Li, Yue Dong, Jiali Wang, Xiuzhu Wang, Yajun Li, Yali Hu, Yayi Hou, Guangfeng Zhao

## Abstract

Macrophages exhibit a high degree of plasticity and play pivotal roles both in the normal physiological cycle of the endometrium and in its regeneration following injury. Although some new subsets of endometrial macrophages have been identified, their origins and functions remain to be further explored. In this study, we employed single-cell sequencing to analyze the endometrium of patients with normal endometrium and intrauterine adhesion (IUA) caused by injury. We identified a unique subset of macrophages distinguished by the expression of GATA6, a marker indicative of cavity macrophages. We verified that these GATA6^+^ macrophages were large peritoneal macrophages (LPMs) that migrated from the peritoneal cavity to the injured endometrium. Upon activation by injured endometrium, these LPMs demonstrated increased expression of Interleukin-33 (IL33), mediated by the Lars-Fos signaling axis, which interacts with the IL33 enhancer. Moreover, our studies revealed that IL33 derived from LPMs inhibited the differentiation of endometrial stromal cells (ESCs) into myofibroblasts, a critical step in the development of endometrial fibrosis. Furthermore, we confirmed the inhibitory effect occurred through the binding of IL33 to the ST2 receptor on ESCs, leading to the upregulation of JMJD3 and subsequent suppression of myofibroblast differentiation. Our findings highlight the essential role of LPMs in promoting endometrial repair and inhibiting fibrosis in IUA.

**Significance Statement:** This study reveals the presence of a unique population of macrophages within the endometrium, originating from large peritoneal macrophages (LPMs), that are pivotal in endometrial repair. This finding offers new insights into the plasticity of macrophages and their potential therapeutic applications in treating endometrial fibrotic disorders.

## Introduction

A healthy woman undergoes approximately 400 menstrual cycles over her lifetime. Under the regulation of hormonal fluctuations, the endometrium sequentially undergoes phases of proliferation, breakdown, and scarless repair. The underlying molecular and cellular mechanisms responsible for this scarless endometrial regeneration remain the focus of intensive research^1^. The human endometrium consists of two distinct layers: the basal layer and the functional layer. It is composed of various cell types, including epithelial cells, stromal fibroblasts, endothelial cells, immune cells, and stem cells^2^. During the process of physiological scarless repair of the endometrium, the controlled activation and regulation of immune cell responses play a pivotal role in orchestrating tissue regeneration^3^. Progesterone withdrawal stimulates endometrial stromal cells to release chemokines, attracting immune cells such as monocytes and macrophages into the endometrial tissue. Subsequently, these monocytes and macrophages rapidly differentiate into a pro-inflammatory phenotype, secreting matrix metalloproteinases (MMPs) to mediate extracellular matrix degradation and tissue breakdown. During the repair phase, these macrophages shift to a pro-reparative phenotype, contributing to tissue remodeling and regeneration^4^. Surgical trauma or severe infections can disrupt the endometrium’s intrinsic repair process, leading to pathological fibrosis and the development of scar tissue.

Intrauterine adhesions (IUA), also known as Asherman Syndrome, stand as a quintessential example of such endometrial fibrotic conditions. Patients with IUA often exhibit irregular menstruation, recurrent miscarriage, and infertility^5, 6^. However, there is still no effective treatment available clinically. The transcervical resection of adhesions (TCRA), which is the preferred clinical approach, fails to fundamentally address the pathological changes of endometrial fibrosis, resulting in a high recurrence rate among IUA patients after surgery^7^. Therefore, understanding the pathophysiological mechanisms driving this condition is crucial to developing more effective and lasting therapeutic strategies.

Macrophages, characterized by their high plasticity, play a pivotal role in both the progression of endometrial repair and fibrosis. The modulation of M1 and M2 macrophage polarization is pivotal in influencing the pathogenesis and progression of IUA^8^. Our preliminary findings revealed a reduction in CD206^+^ M2 macrophages in the endometrium of patients with IUA^9^. However, the M1/M2 classification of macrophages is now widely regarded as insufficient for accurately delineating the functional diversity and phenotypic heterogeneity of macrophages in specific biological contexts. Currently, an increasing number of novel macrophage subsets in the endometrium have been discovered to be involved in endometrial fibrosis. Our previous research found that CD301^+^ macrophages constitute a unique subset within the endometrial microenvironment. These macrophages facilitate the transdifferentiation of ESCs into myofibroblasts, driving excessive extracellular matrix deposition and thereby playing a key role in the pathogenesis of endometrial fibrosis^10^. Additionally, we also discovered that IL34 can promote the differentiation of CX3CR1^+^ macrophages in the endometrium, thereby exacerbating the severity of IUA^11^. This underscores the distinct roles played by different subsets of endometrial macrophages. Moreover, it is widely acknowledged that endometrial macrophages consist of both tissue-resident macrophages and macrophages or monocytes recruited from the periphery that differentiate locally, with each potentially exhibiting unique functions depending on their origin.

In this study, we identified a distinctive subset of macrophages originating from the peritoneal cavity, characterized by the expression of GATA6 in the injured endometrium of IUA patients, based on 10x Genomics single-cell RNA sequencing (scRNA-seq). GATA6 and CD102 are the markers of LPMs. Our findings revealed that the depletion of LPMs exacerbated the progression of injury-induced endometrial inflammation and fibrosis in mice. Upon activation by injured endometrium, these LPMs demonstrated increased expression of Interleukin-33 (IL33). Notably, LPMs-derived IL33 inhibited the differentiation of ESCs into myofibroblasts, thereby mitigating endometrial fibrosis through the ST2-JMJD3 pathway. Mechanically, Lars enhanced IL33 expression by promoting Fos binding to the IL33 enhancer.

Collectively, our research elucidates the functional role and molecular mechanism of LPMs in endometrial fibrosis and suggests that targeting LPMs-derived IL33 may represent a promising therapeutic approach for treating IUA.

## Methods

### Human samples

The human endometrial samples and procedures in this study were conducted with approval from the Ethics Committee of Nanjing Drum Tower Hospital. All participants provided written informed consent. Endometrial samples were obtained from women aged 22 to 40 years during the late proliferative phase of their menstrual cycle, collected during hysteroscopic examination. Samples were sourced from patients with IUA caused by dilation and curettage (D&C), whose endometrial condition scored above 8 according to the American Fertility Society’s criteria, and from non-IUA patients who presented with tubal infertility, normal ovarian function, typical menstrual blood volume, and an endometrial thickness of at least 7 mm just before ovulation. Women were excluded from the study if they had any of the following conditions: positive serological results for HIV, hepatitis B or C, or syphilis; a history of tuberculosis; chronic endometritis; or vaginal bacterial or fungal infections. Additionally, uterine malformations were ruled out via ultrasonography before sample collection.

### Single-cell RNA-seq data processing

The methodology employed in this study is consistent with our previous reporte^10^. In brief, the endometrial samples were rinsed with PBS, cut into small pieces, and then digested with 0.1% trypsin for 8 minutes, followed treatment with 0.8 mg/ml Collagenase Type I for 60 minutes at 37°C with 5% CO_2_. The released cells were filtered, centrifuged, and treated with red blood cell lysis buffer before resuspension in PBS for single-cell 3’ cDNA library preparation. Cells expressing fewer than 200 genes or with mitochondrial gene content > 15% were excluded. Single-cell encapsulation, cDNA library synthesis, and RNA sequencing were performed by Gene Denovo. The data were aligned to the human genome (GRCh38) using the STAR algorithm, and the UMI count matrix was processed using the Seurat toolkit for normalization and log-transformation.

### Cell culture

LPMs and ESCs were obtained from female Balb/C mice (8-10 Weeks) as previously described^12, 13^. The peritoneal cavity of mice was lavaged with 5 mL of cool PBS for 2 min in a sterile environment. Isolated peritoneal cells were cultured in DMEM (Giboc, Grand Island, NY, USA) containing 10% fetal bovine serum (FBS; Gibco, USA), 100 U/mL penicillin, and 100 μg/mL streptomycin (100 µg/mL; Gibco, Grand Island, NY, USA) and cultured at 37 °C with 5% CO_2_ and saturated humidity. After 4h, the cells were rinsed 3 times with warm PBS to eliminate non-adherent cells, leaving only the adherent cells, which were identified as LPMs. In sterile environment, the uterus of mice was digested by mixture combine with DF-12 (Giboc, Grand Island, NY, USA), collagenase type I (Biosharp, CN), and DNase (Roche, Switzerland). Then 40μM cell strainers were used to depart single-cells from uterus tissue. The ESCs were cultured in DF-12 containing 10% fetal bovine serum (FBS; Gibco, USA), 100 U/mL penicillin, and 100 μg/mL streptomycin (100 µg/mL; Gibco, Grand Island, NY, USA) and cultured at 37 °C with 5% CO_2_ and saturated humidity. ESCs were used in P1-P3 in all experiments.

### Animals and experimental protocol in vivo

Female Balb/C, C57BL/6 or CD45.1 mouse weighting 20-23g (8-10 Weeks) were purchased from Nanjing Cavans Biotechnology Co, Ltd. They were housed in specific pathogen-free (SPF) condition with 12 h dark/light cycle, and could free access standard chow diets and water. All animal experiments were carried out in accordance with the guidelines of the Experimental Animals Management Committee (Jiangsu Province, China) and were approved by the Ethics Review Board for Animal Studies of Nanjing Drum Tower Hospital.

### Electric tool-scratching mouse endometrial injury model

The electric tool-scratching mouse endometrial injury model was like the previous reported^14^. In brief, after anesthetizing the mice with isoflurane, the probe with a rough surface was insert into uterus and push the vibrator switch for 3-cycle (10s shake and 2s pause). This can cause mechanical damage to the endometrium. This method couldn’t damage the integrity of the peritoneum, ensuring that there was no loss of LPMs. We detected the inflammation stage in 6 h after the model established, and the fibrosis stage in 7 Days after the model established.

### LPMs depletion

After anesthetizing the mice with isoflurane, the mouse was lavage 2 times with PBS in a sterile environment. Endometrial injury model was established in mice after 24h.

### CD45.1^+^/CD45.2^+^ LPMs adoptive transfer

100uL clodronate liposomes (CLLs) were intraperitoneally injected into WT mice to deplete CD45.2^+^LPMs. After 48h, CD45.1^+^LPMs were obtained from CD45.1 mouse and adoptive injected to WT mice. Endometrial injury model was established in mice after 24h after successfully transfer^15^.

### Mechanical injury induces ESCs apoptosis

After the ESCs were affixed to the cell dishes for 24h, they were forcefully detached from the dishes with DMEM complete medium and strongly blown up for 50 times. The supernatants were collected after 12h of culturing the damaged ESCs and the mechanical damage-induced AES was obtained after filtration with a 0.22 μM filter.

### Cell migration

LPMs were seeded in 3uM transwells, while apoptosis ESCs were seeded in the lower compartment of the transwells. The transwells were then put into 4% PFA for 15min in room temperature and subsequently washed 3 times with PBS. Following this, the transwells were placed in 1% ammonium oxalate crystal violet for 45min in room temperature and washed 3 times with PBS. Any LPMs that had not migrated to the opposite side were carefully removed using a cotton swab. The transwells were then placed under a microscope for observation. Three fields of view were randomly selected for cell counting, and the average value was counted.

### RNA isolation and quantitative real-time PCR (qPCR)

Following the protocol of manufacturer, total RNA was extracted by Trizol regent (Vazyme Biotech, China). Total RNA was reversed to cDNA by HiScript ll Q RT SuperMix for qPCR (Catalogue # R222-01; Vazyme Biotech, China). The method of 2^-△△CT^ was used to analysis the target genes expression levels, normalized by β-actin expression. The primers used in this study are listed in Supplementary Table 1.

### Western blot analysis (WB)

The protein of cells was lysed by RIPA lysis buffer (Beyotime Biotechnology) with protease inhibitor cocktail and phosphatase inhibitor cocktail (Roche), and clarified by centrifugation at 12,000 × g for 20 min. The BCA protein assay kit (Thermo) was used to determine the concentration of protein samples. Then protein samples were boiled in 99[ in Dry Bath Incubator, mixing 5X loading buffer. Protein samples were separated using SDS-PAGE, transferred to PVDF membranes and incubated in 5% bovine serum albumin (BSA) in room temperature for 2h. Primary antibody dilutions were added and incubated overnight at 4°C in a shaker. The membrane was then incubated with secondary antibody coupled with HRP for 2 h at room temperature.

Protein signals were observed using ECL solution and quantitative analysis of gray values was performed using Image J. The antibodies used in this study are listed in Supplementary Table 2.

### Gene silencing of *Lars*, *Trem1* and *Il33* using siRNA

Cells were transfected using Lipofectamine 3000 according to the manufacturer’s guidelines (Invitrogen, USA). Si-*Il33* was designed and purchased in Ribobio (Guangzhou, China). Si-*Lars* and si-*Trem1* were designed and purchased in Genepharma (Shanghai, China). The sequences of siRNA are listed in Supplementary Table 3.

### Immunofluorescence

The cells were culture in coverslip and subsequently fixed in 4% paraformaldehyde (PFA) and wash 3 times using PBS. After permeabilization with 0.2% Triton X-100, cells were incubated in 1% BSA in room temperature for 1h. Primary antibody dilutions were added and incubated overnight at 4°C. After that, the cells were incubated with secondary antibody coupled with fluorescence for 2 h at room temperature in dark. The nuclei were stained by 4′,6-diamidino-2-phenylindole (DAPI) and observed under FV3000 Laser Scanning Confocal Microscope (Olympus Corporation, Tokyo, Japan). Analyzing mean optical density or pearson’s coefficient with image J. The antibodies used in this study are listed in Supplementary Table 2.

### Flow cytometry

The labeling of LPMs with fluorescent beads (Fluoresbrite® YG Microspheres, Polysciences Inc.) was performed using a method similar to that previously reported^16^. In brief, before establishing endometrial injury model in mouse, we injected a 100uL mixture (10uL fluo Bead and 90uL PBS) to label the LPMs. After establishing endometrial injury model for 6h, we acquired the peritoneal lavage and uterine single-cell suspension. Fluorescent antibodies were used to label cells, and the antibodies used in this study are listed in Supplementary Table 2. All flow cytometry data were acquired using the Beckman Coulter Cytoflex S (Beckman Coulter, CA, USA) or BD FACS Calibur cytometer (BD Biosciences, San Diego, CA, USA) and subsequently analyzed using FlowJo software (Treestar, Inc., San Carlos, CA, USA).

### Immunohistochemistry

Paraffin blocks were sectioned into 2-μm-thick slices and mounted onto glass slides. To block endogenous peroxidase activity, the sections were treated with 3% H_2_O_2_, and antigen retrieval was performed using Immunohistochemistry Universal Antigen Recovery Solution. The sections were then incubated overnight at 4°C with diluted primary antibodies. Subsequently, they were incubated with HRP-conjugated secondary antibody, and the antigen signal was visualized with DAB. The sections were counterstained with hematoxylin, sealed with neutral resin, and the positive reactants were visualized under a microscope. The immunohistochemical staining was quantified by calculating the mean optical density using Image software. The antibodies used in this study are listed in Supplementary Table 2.

### Bulk RNA-sequencing and analysis

LPMs were treated with or without ASE for 6h, and total RNA was extracted by Trizol regent. Transcriptome sequencing analysis was performed on an Illumina NovaSeq6000 by Gene Denovo Biotechnology Co. (Guangzhou, China). Differential expression analysis between two different groups was performed with DESeq2 software. Genes with a false discovery rate (FDR) below 0.05 and absolute fold change ≥ 1.5 were considered differentially expressed genes (DEGs). KEGG which was the major public pathway-related database for pathway enrichment analysis was performed for all DEGs. It identified significantly enriched metabolic pathways or signal transduction pathways in DEGs comparing with the whole genome background.

### Dual luciferase reporter assay

We selected a 2,000 bp sequence upstream of the transcription start site (TSS) of the *Il33* gene (gene ID: 77125) as the promoter region (>NC_000085.7:29925060-29927060 Mus musculus strain C57BL/6J chromosome 19, GRCm39). The enhancer region for Il33 was chosen from the NCBI database (>NC_000085.7:29922546-29923676 Mus musculus strain C57BL/6J chromosome 19, GRCm39). Corresponding luciferase reporter plasmids were constructed for both regions. Using the JASPAR database, we predicted the top three FOS::JUN binding sites and deleted these binding sites to generate the corresponding mutant plasmids. Transfection was performed using Lipofectamine 3000 (Invitrogen), according to the manufacturer’s protocol. For reporter assays, cells seeded in 24-well plates were transfected with 800 ng IL33-promoter-WT or IL33-promoter-MUT or IL33-enhencer-WT or IL33-enhencer-MUT, 800ng Fos, and 100 ng Renilla plasmid. After 48 hours, the cells were lysed, and reporter activity was measured using the Dual-Luciferase Reporter Assay System (Vazyme Biotech Co., Ltd.) following the manufacturer’s instructions.

### Statistical analysis

All the experiments were randomized and blinded. All the experiments were repeated at least three independent times. All the values presented in the graphs are shown as the means ± SEMs. The Shapiro-Wilk test was used to confirm whether the data were normally distributed. One-way ANOVA was used to analyze the data among more than two groups, and unpair Student’s t tests were used to compare two groups. GraphPad Prism 5 Demo (GraphPad Software Inc., La Jolla, CA, USA) was utilized for statistical analysis.

## Results

### Result 1: GATA6^+^ macrophages in the injured endometrium are derived from the peritoneal cavity

By conducting 10x Genomic single-cell sequencing analysis on the endometrium samples collected from three healthy individuals and three patients with IUA, we discovered a distinct subpopulation of macrophages that express the cavity macrophage marker GATA6. This particular subpopulation is enriched in the endometrium of patients with IUA (Fig.1A and B). The immunostaining further showed that GATA6 was co-stained with macrophages marker CD68 in IUA endometria (Fig.1C). Since the uterus is located in the peritoneal cavity, we hypothesize that this population of GATA6^+^ macrophages may originate from the peritoneal cavity. To validate this hypothesis, we established a mouse endometrial injury model. Six hours after inducing the injury, we collected peritoneal lavage fluid and uterine single-cell suspensions for flow cytometry analysis, following the gating strategy outlined in Supplementary fig. 1A-B. The results revealed that endometrial injury triggered inflammation in the uterus, marked by a notable surge in the number of CD11b^+^ cells (Fig.1D). Furthermore, there was a significant decline in LPMs in the peritoneal cavity (Fig.1E) and a concomitant rise in LPMs in the uterus post-injury (Fig.1F), indicating that LPMs may migrate from the peritoneal cavity to the injured uterus. To further confirm this phenomenon, we conducted CD45.1^+^/CD45.2^+^ LPMs adoptive transfer experiments^17^ (Fig.1G). The flow cytometry results showed that CD45.1^+^ cells were significantly reduced in the peritoneal cavity of Injured group (6h group) compared to Control group (Supplementary fig. 1C). Conversely, the proportion of CD45.1^+^ cells was significantly elevated in the damaged uterus, and these CD45.1^+^ cells were identified as F4/80^+^CD102^+^cells (LPMs). Meanwhile, CD45.2^+^ cells in the damaged uterus were F4/80^+^CD102^-^ cells (Fig.1H). These findings strongly indicated that LPMs migrate from the peritoneal cavity to the damaged uterus both in human and mouse.

**Fig 1.**
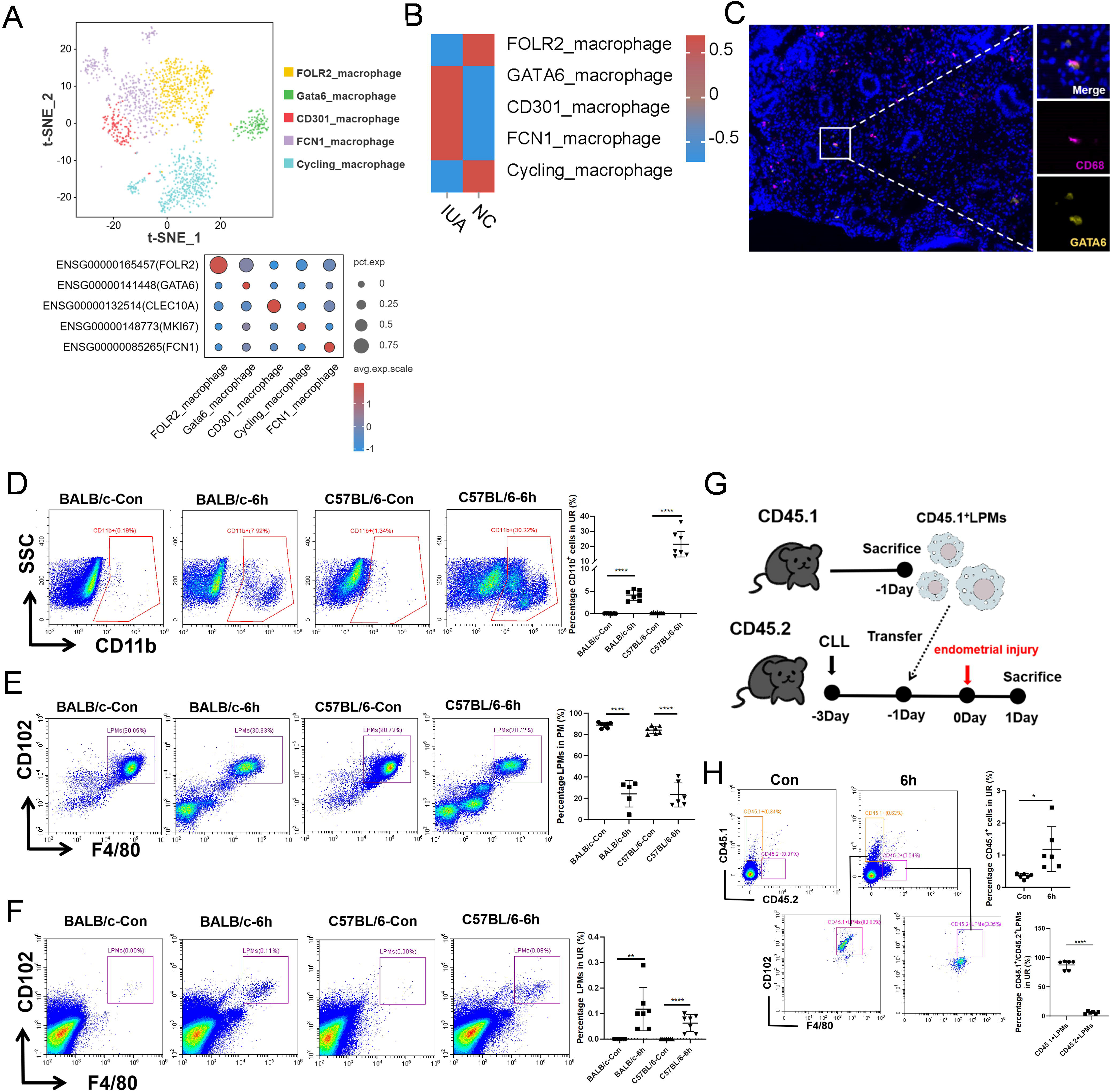
GATA6^+^ macrophages (LPMs) in human or mouse endometrium between IUA patients and healthy controls. A. The tSNE plot displaying macrophages from normal control and patients with IUA (n = 3). B. Differences in macrophage subpopulations between normal control and patients. C. Representative immunofluorescence images of GATA6 and CD68 from samples of patients with IUA. D. Flow cytometry analysis of changes in the proportion of CD11b^+^ cells in the uterus of normal control and endometrial injury mice (n=6). E. Flow cytometry analysis of changes in the proportion of LPMs in the peritoneal cavity of normal control and endometrial injury mice (n=5-6). F. Flow cytometry analysis of changes in the proportion of LPMs in the uterus of normal control and endometrial injury mice (n=6). G. Schematic diagram of the flow of CD45.1/CD45.2 adoptive transfer experiments. H. Flow cytometry analysis of changes in the proportion of CD45.1^+^ in the uterus of normal control and endometrial injury mice (n=5-6). Values are mean±SD. *p<0.05, **p<0.01, ***p<0.001, ****p<0.0001, ns denotes p>0.05 (by unpaired Student’s t test or one-way ANOVA)

### Result 2: Depletion of LPMs exacerbates inflammation and fibrosis within the injured endometrium

To investigate the role of LPMs in damaged endometrium, we initially employed intraperitoneal lavage (PL) to specifically remove LPMs from the peritoneal cavity. The results showed that PL did not elicit a systemic inflammatory response or uterine damage in mice (Supplementary fig. 2A, B, C). Flow cytometry analysis further confirmed that PL effectively and stably depleted LPMs in the peritoneal cavity of mice (Fig.2A). After establishing the endometrial injury model, we observed a notable absence of significant LPMs cluster in the damaged uterus of the PL group (Fig. 2B), suggesting that LPMs depletion prevented their migration to the injured uterus. Our results also demonstrated that the injury significantly elevated the systemic inflammation level in mice, as evidenced by the marked upregulation of serum inflammatory factors IL-1β, IL-6, and TNF-α. This inflammatory response was further exacerbated by the depletion of LPMs (Fig.2C). Additionally, following endometrial injury, we observed a significant reduction in the number of endometrial glands, an increase in collagen deposition, an upregulation of the fibrosis marker α-SMA, and a decrease in the expression of the cell proliferation marker Ki67. Depletion of LPMs exacerbated these injury and fibrotic symptoms (Fig.2D-G). Taken together, these data suggested that LPMs migrating into damaged endometrium might play a protective role, promoting the repair of endometrial injury.

**Fig 2.**
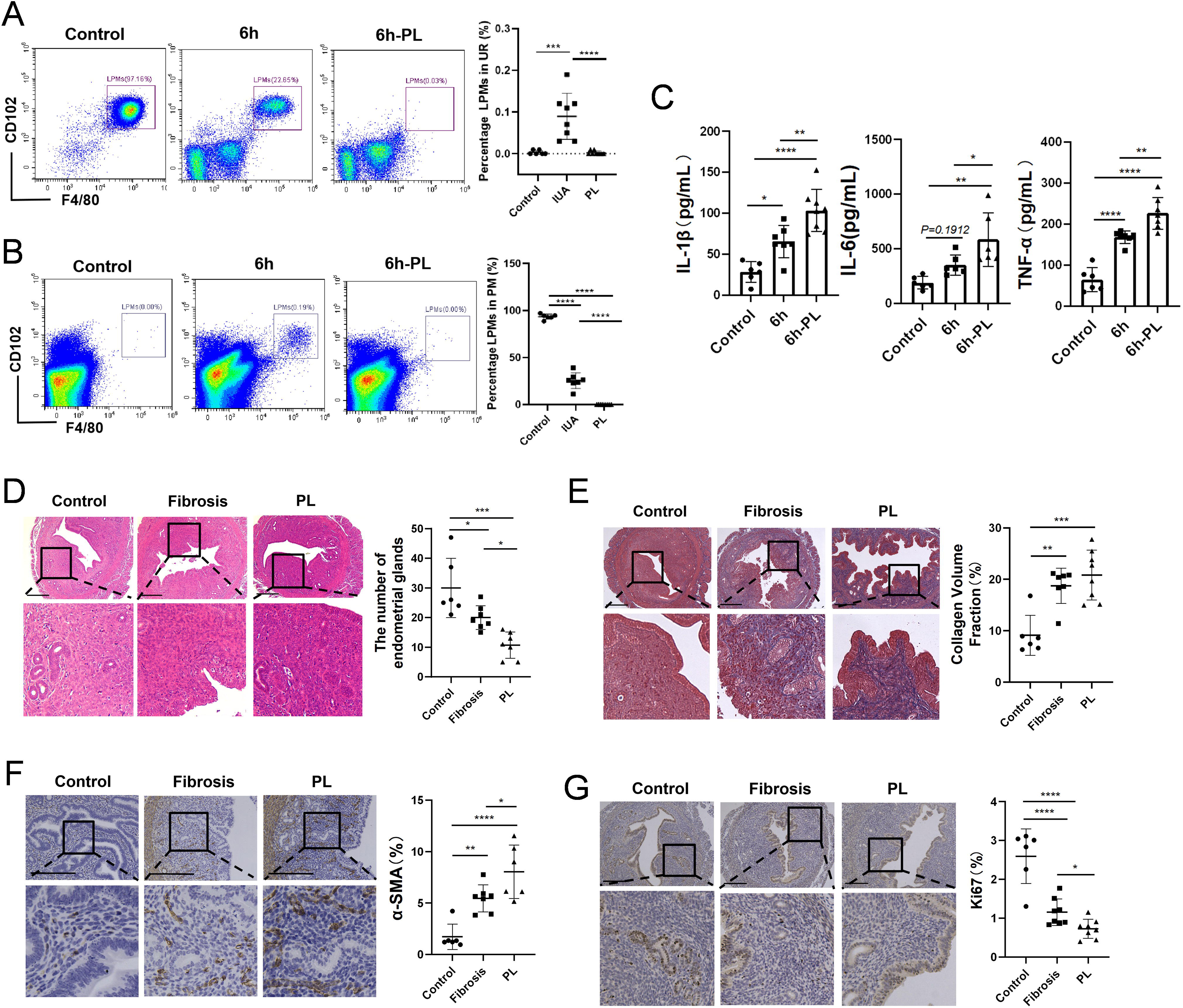
Depletion of LPMs exacerbates inflammation levels and endometrial fibrosis in endometrial injury mice. A. Flow cytometry analysis of changes in the proportion of LPMs in the peritoneal cavity (n=6), PL means depleting LPMs by intraperitoneal lavage. B. Flow cytometry analysis of changes in the proportion of LPMs in the uterus (n=6). C. Serum concentrations of IL-1β, TNF-α, and IL-6 were measured by ELISA (n=6). D. HE staining of endometrial tissues obtained from mice (n=6-7). E. Masson’s trichrome staining of endometrial tissues obtained from mice (n=6-7). F. Representative images of IHC for α-SMA staining in the mice endometria (n=6-7). G. Representative images of IHC for Ki67 staining in the mice endometria (n=6-8). Scale bar indicates 200 μm. Values are mean±SD. *p<0.05, **p<0.01, ***p<0.001, ****p<0.0001 ns denotes p>0.05 (by one-way ANOVA).

### Result 3: The LPMs entering the damaged endometrium undergoes phenotypic changes

We utilized fluorescent beads to track LPMs (Beads^+^LPMs). These beads specifically labeled LPMs with a 91% efficiency (Supplementary fig. 3A) and did not stain the undamaged uterus with fluorescence from the beads (FITC) (Supplementary fig. 3B). We then collected peritoneal lavage and uterine single-cell suspensions at 0, 6, 12, 24, and 48 hours after establishing the endometrial injury model, and analyzed these samples using flow cytometry. The results revealed that Beads^+^LPMs persistently migrated into the damaged uterus within 48 hours, accompanied by a decrease in the LPMs marker CD102 (Fig.3A). We further confirmed the labeling of LPMs by immunofluorescence co-localization, which showed that Beads and CD102 co-localized at 6 and 12 hours (Fig.3B). Additionally, there was no co-localization of Bead fluorescence and Tunel fluorescence at 24 and 48 hours, indicating that the labeled LPMs did not undergo apoptosis at these time points (Fig.3C). These findings indicated that the observed CD102 reduction within Bead^+^ cells were reflective of alterations in LPMs specifically, rather than changes in other macrophage populations present in the uterus. The Beads^+^CD11b^+^ cells in the peritoneal cavity exhibited a similar trend of change as LPMs, further validating the success of our LPMs labeling (Supplementary fig. 3C, D). We also observed that LPMs could penetrate the uterine muscle layer and accumulate at the damaged endometrium, indicating that LPMs alleviate endometrial fibrosis by interacting with cells at the endometrial site (Fig.3D).

**Fig 3.**
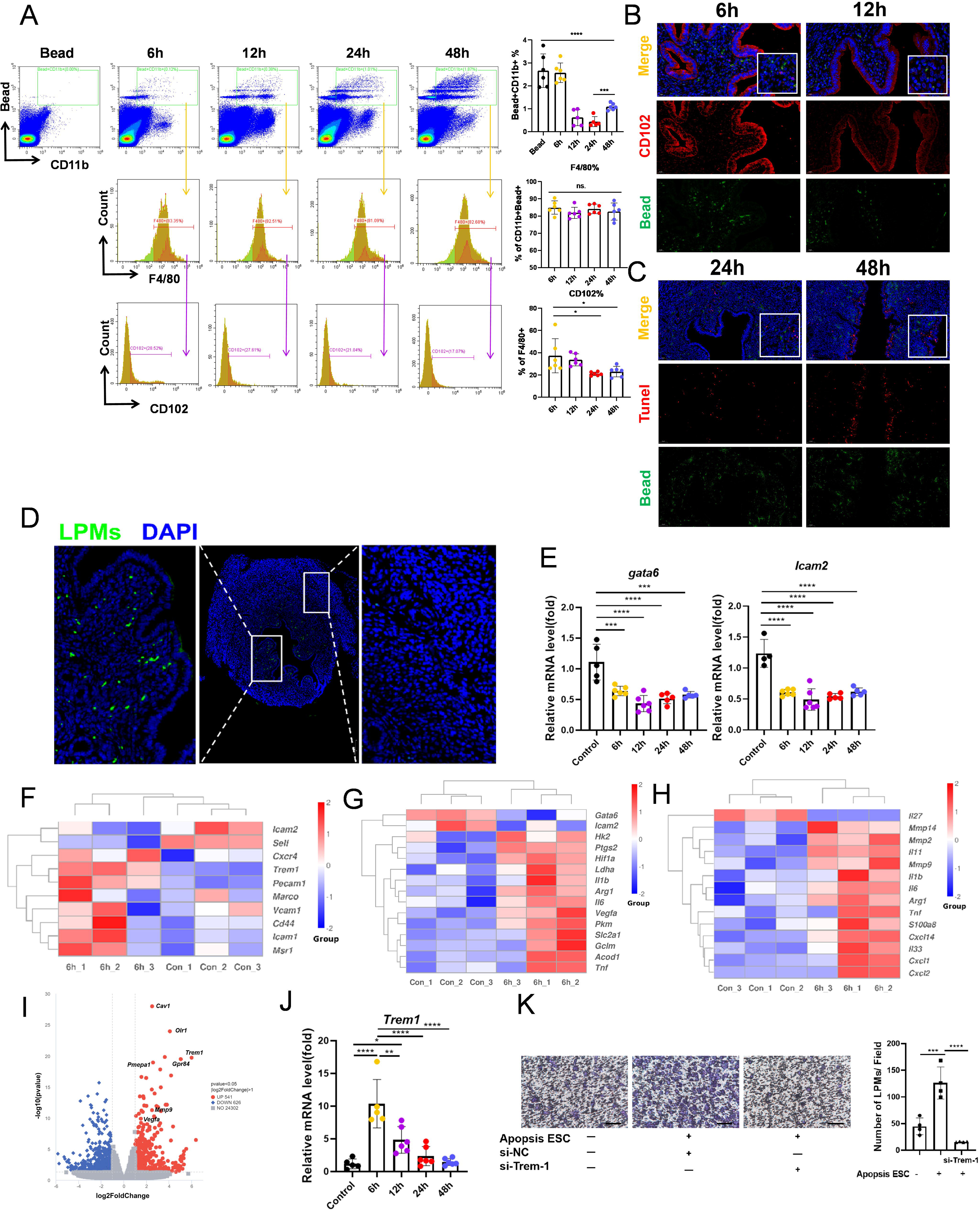
LPMs responding to wound signaling change their transcriptome profile. A. Flow cytometry analysis of changes in the proportion of Beads^+^LPMs and the change of them in the mice uterus (n=5-6). B. Representative images of immunofluorescence for CD102 and Bead in the endometrial injury mouse endometria. C. Representative images of immunofluorescence for tunel and Bead in the endometrial injury mouse endometria. D. Representative images of immunofluorescence for Bead in the endometrial injury mouse uterus. E. The gene levels of *Gata6* and *Icam2* were detected with qRT-PCR (n=5-6). F. The thermogram of migration-associated molecule related genes expression pattern. G. The thermogram of early-phase wound macrophages and LPMs phenotype related genes expression pattern. H. The thermogram of cytokines related genes expression pattern. I. The volcano figure showed foldchanges of genes in LPMs incubated with control or AES-treated LPMs (6h). J. The gene levels of *Trem1*were detected with qRT-PCR (n=5-6). K. Crystal violet staining of LPMs cocultured with apopsis ESCs or not. (scale bar: 200 μm) Values are mean±SD. *p<0.05, **p<0.01, ***p<0.001, ****p<0.0001, ns denotes p>0.05 (by unpaired Student’s t test or one-way ANOVA)

Endometrial injury can result in some cell death of stromal cells in uteri. Therefore, we established a method to induce the death of endometrial stromal cells (ESCs) through mechanical force (Supplementary fig. 4A). Subsequently, we treated LPMs with the apoptotic ESCs supernatant (AES). Consistent with in vivo findings, the LPMs markers *Gata6* and *Icam2* (encoding CD102) exhibited significant downregulation starting from 6 hours (Fig. 3E). We also subjected LPMs to AES for 0 or 6 hours for comprehensive genome-wide mRNA expression analysis. Principal Component Analysis (PCA) of the samples revealed distinct clustered distributions between the two groups, which could be clearly separated into two sample clusters (Supplementary fig. 4B). Compared to the control group, a total of 1,187 differentially expressed genes (DEGs) were identified in the LPMs treated with AES for 6 hours, with 541 genes up-regulated and 646 genes down-regulated (Fig. 3I). Heatmaps of DEGs indicated that gene expression patterns clustered distinctly after unsupervised clustering (Supplementary fig. 4C). Migration-associated molecules (Fig. 3F), early-phase wound macrophage-associated molecules and phenotypic markers of LPMs (Fig. 3G; Supplementary fig. 4D), and cytokines (Fig. 3H) displayed different transcriptomic profiles, suggesting alterations in AES-activated LPMs compared to their resting state. The volcano plot of differential genes highlighted that Triggering receptor expressed on myeloid cells 1 (Trem1) was significantly overexpressed at 6 hours (Fig. 3I). Trem1, a cell surface receptor found on monocytes or tissue-resident macrophages, can be activated by damage-associated molecular patterns (DAMPs) and is transcriptionally regulated by Hif1α^18^. The qRT-PCR results verified that *Trem1* expression was significantly high at 6 hours, followed by a decline and a gradual return to baseline levels (Fig. 3J). Trem1 is known to promote macrophages migration and adhesion^19^. Expectedly, Results from the transwell migration assay indicated that AES treatment enhanced LPMs migration, while knockdown of *Trem1* inhibited this migration process (Fig. 3K). These findings implied that DAMPs released from the damaged endometrium may promote LPMs migration by upregulating *Trem1* expression. These findings implied that DAMPs released from the damaged endometrium may promote LPMs migration by upregulating *Trem1* expression.

What’s interesting was that the DAMPs could also change the LPMs phenotype. However, phenotypic changes in LPMs related to their functional alterations and the repair of the injured endometrium remains unclear.

### Result 4: Changed LPMs phenotypically secrete IL33 to promote endometrial repair

To further determine the proteins integral to macrophages functionality secreted by activated LPMs, the gene expressions of secretory proteins transcriptomic profiles (Fig. 3H) were investigated by qRT-PCR. Notably, *Il33* and *Arg1* were significantly upregulated in LPMs within the 6h group (Supplementary Fig. 5A). ScRNA-seq analysis also revealed the specific and high expression of *IL33* in LPMs (Fig. 4A), with co-localization of Il33 and Gata6 evident (Fig. 4B; Supplementary Fig. 5B).

**Fig 4.**
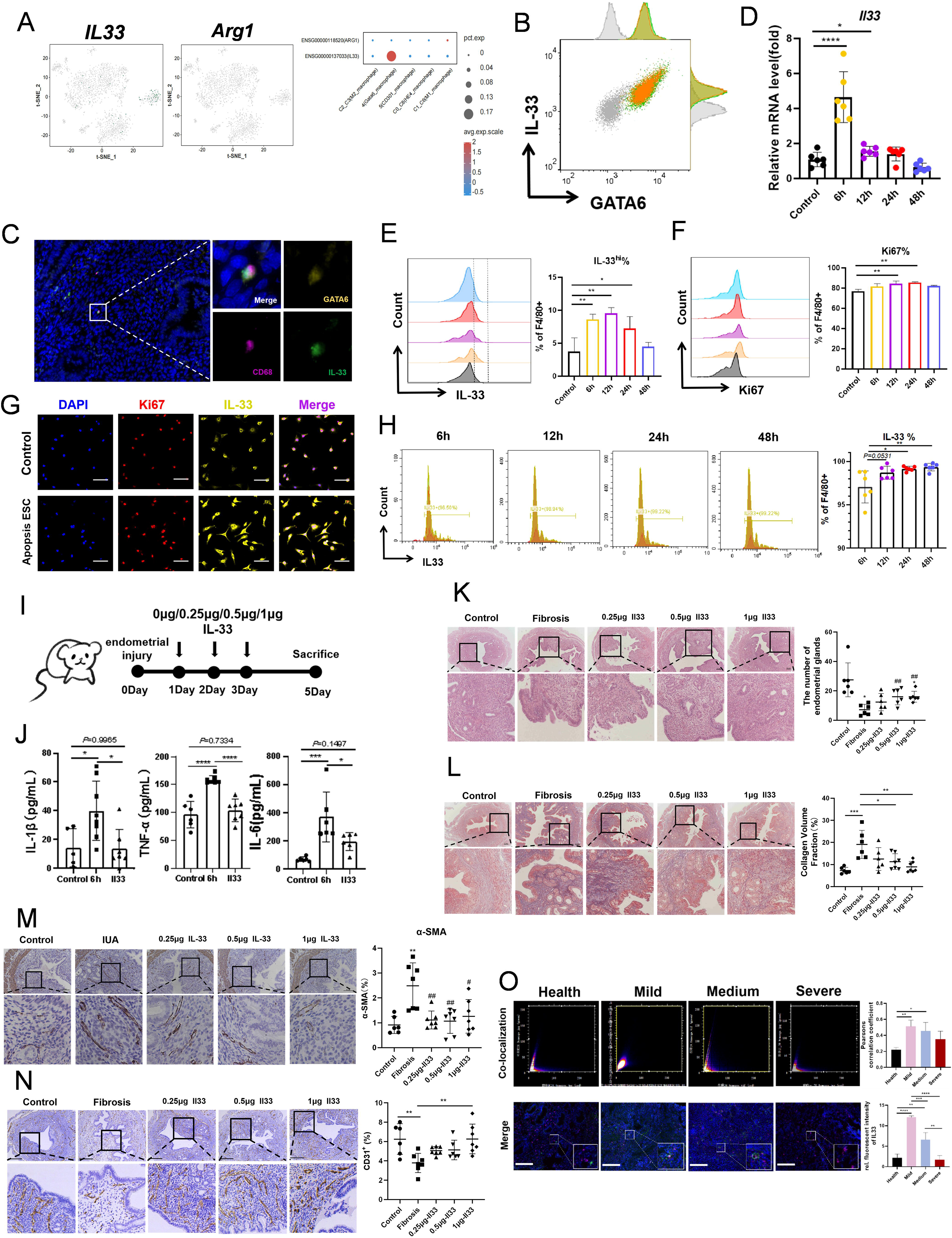
Activated LPMs highly express Il33 to relives endometrial injury. A. The tSNE plot showing the expression of *Il33* and *Arg1* for macrophages. B. Flow cytometry analysis of GATA6 and Il33 in LPMs. C. Representative immunostaining images of CD68, GATA6 and IL33 from samples of IUA patients. D. The gene levels of *Il33* was detected with qRT-PCR (n=6). E. Flow cytometry analysis of Il33 in AES treated LPMs (n=4). F. Flow cytometry analysis of Ki67 in AES treated LPMs (n=4). G. Representative immunofluorescence images of Il33 and Ki67 from samples of LPMs (scale bar: 40 μm). H. Flow cytometry analysis of changes in the proportion of Bead^+^CD11b^+^IL33^+^ cells in the endometrial injury mice uterus (n=5-6). I. Schematic diagram of the procession of endometrial injury model. J. Serum concentrations of IL-1β, TNF-α, and IL-6 were measured by ELISA (n=7). K. HE staining of endometrial tissues obtained from mice (n=7). L. Masson’s trichrome staining of endometrial tissues obtained from mice (n=7). M. Representative images of IHC for α-SMA staining in the mice endometria (n=7). N. Representative images of IHC for CD31 staining in the mice endometria (n=7). O. Representative immunofluorescence images of GATA6 and CD68 from samples of patients with IUA (n = 3-5). Values are mean±SD. *p<0.05, **p<0.01, ***p<0.001, ****p<0.0001, ns denotes p>0.05 (by unpaired Student’s t test or one-way ANOVA)

Importantly, we identified CD68^+^GATA6^+^IL33^+^macrophages in the endometrium of IUA patients (Fig. 4C). At 6 hours of AES treatment, LPMs exhibited peak *Il33* gene expression (Fig. 4D), with IL33 protein levels reaching their maximum at 12 hours (Fig. 4E; Supplementary Fig. 5C), suggesting that DAMPs released from injured ESCs may activate LPMs, leading to the upregulation of IL33. To confirm our hypothesis, we treated LPMs with ATP, a classical DAMPs. The results showed that ATP promoted IL33 expression at both the gene and protein levels (Supplementary Fig. 5D, E, F). Concurrently, AES treatment significantly upregulated Ki67 expression in LPMs (Fig. 4F, G). A marked increase in the proportion of CD11b^+^F4/80^+^IL33^+^ cells was also observed during the progression of endometrial injury in vivo (Fig. 4H). Altogether, damaged endometrial tissue can activate LPMs, resulting in the robust expression of IL33.

To determine whether the inhibitory effect of LPMs on endometrial fibrosis is mediated through the secretion of IL33, we directly treated the endometrial injury mouse model with IL33 recombinant protein. Following the intraperitoneal administration of 0.25 μg, 0.5 μg, or 1 μg of IL33 recombinant protein, we evaluated the condition of mice with endometrial injury (Fig. 4I). The results showed that intraperitoneal injection of IL33 inhibited effectively both uterine inflammation and systemic inflammation induced by endometrial injury (Supplementary fig. 5G, Fig. 4J). Additionally, it triggered a type 2 immune response that facilitated tissue repair (Supplementary fig. 5H). Importantly, this treatment did not increase serum levels of Il33 and sST2, a specific soluble receptor for IL33 (Supplementary fig. 5I). Il33 was found to preserve the normal morphology and quantity of uterine glands (Fig. 4K), inhibit collagen deposition (Fig. 4L) and α-SMA expression in the endometrium (Fig. 4M), and enhance CD31 expression (Fig. 4N). These results suggested that intraperitoneal injection of Il33 can alleviate injury induced endometrial fibrosis.

Furthermore, we collected endometrial tissue samples from patients with IUA of varying degrees of clinical severity. Our analysis revealed a notable trend: the Pearson’s correlation coefficient between CD68 and IL33 was higher in patients with mild disease, and this correlation progressively diminished as the severity of the disease increased. Consistent with this finding, the fluorescence intensity of IL33 was also observed to be higher in patients with less severe disease, decreasing as the disease severity escalated (Fig. 4O). In brief, LPMs migrated to the damaged endometrium and secreted IL33 to promote uterine repair and inhibit endometrial fibrosis.

### Result 5: IL33 inhibits the differentiation of ESCs into myofibroblasts by upregulating JMJD3 via ST2

ST2 is the sole receptor for IL33 signaling. Our observations revealed a notable upregulation of ST2 expression in the uterine tissues of mice with endometrial injury, with specific localization on ESCs (Fig. 5A, B). A key factor in the development of endometrial fibrosis is the accumulation of myofibroblasts, with the transdifferentiation of endometrial stromal cells (ESCs) into myofibroblasts^10, 20^. We induced ESCs differentiation toward myofibroblasts using TGF-β1 and subsequently co-cultured them with non-activated LPMs, activated LPMs, activated LPMs with *Il33*-knockdown, or recombinant IL33 (Fig. 5C). The results showed that TGF-β1 led to a significant increase in α-SMA expression in ESCs, which was not substantially affected by non-activated LPMs. However, activated LPMs inhibited α-SMA expression, and this suppressive effect was abrogated when *Il33* was knocked down in the activated LPMs. Additionally, activated LPMs suppressed the gene expression of *Acta2* (encoding α-SMA) and *Col1a1* (encoding Collagen I), but this inhibition was reversed when *Il33 was* knocked down prior to LPMs activation (Fig. 5D). Similar results were obtained upon the addition of recombinant IL33 protein (Supplementary Fig. 6A, B, C). Furthermore, our findings revealed that activated LPMs elevated both the mRNA and protein levels of *Kdm6b* (encoding JMJD3) in ESCs, and this increase was reversed when *Il33* was knocked down (Fig. 5E, F). Prior research has shown that JMJD3 in renal interstitial cells exhibits anti-fibrotic effects by limiting multiple pro-fibrotic signaling pathways^21^. These results indicate that IL33 inhibits TGF-β1-induced fibrosis by upregulating JMJD3 expression in ESCs.

**Fig 5.**
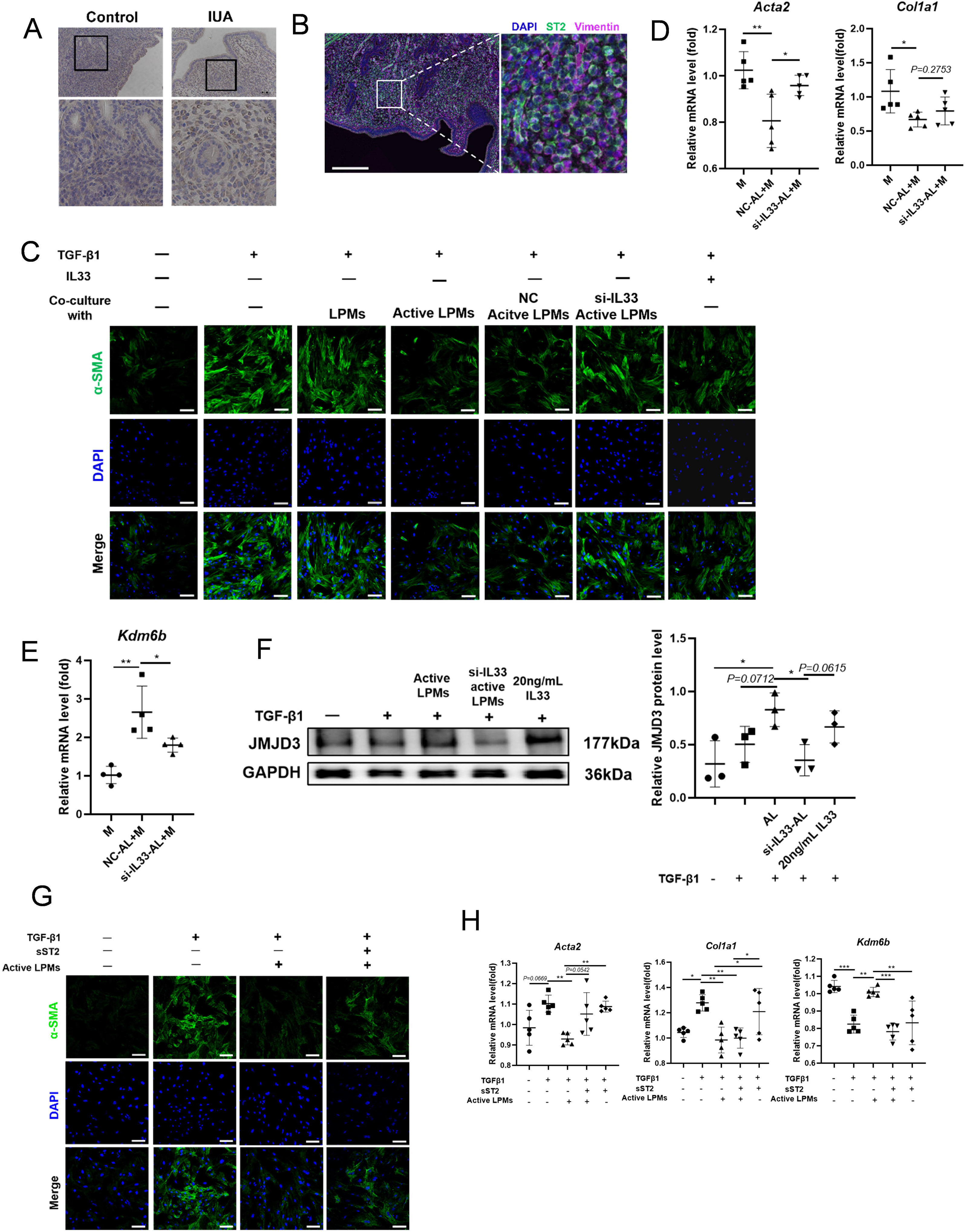
LPMs-derived IL33 inhibits differentiation of ESCs into myofibroblasts via JMJD3. A. Representative images of IHC for ST2 staining in the mice endometria (n=3). B. Representative images of immunofluorescence for ST2 and Vimentin in the endometrial injury mouse endometria (n=3). C. Representative immunofluorescence images of α-SMA from samples of ESCs (n=3) (scale bar: 100 μm). (with or without TGFβ1(10ng/mL), with or without Il33 (20ng/mL), with or without active-LPMs/ LPMs in resting) D. The gene levels of *Acta2* and *Col1a1* were detected with qRT-PCR (n=5). E. The gene levels of *Kdm6b* were detected with qRT-PCR (n=4). F. The protein levels of Jmjd3 were detected with WB (n=3). G. Representative immunofluorescence images of α-SMA from samples of ESCs (n=3) (scale bar: 100 μm). F. The gene levels of *Acta2, Col1a1* and *Kdm6b* were detected with qRT-PCR (n=5). Values are mean±SD. *p<0.05, **p<0.01, ***p<0.001, ****p<0.0001, ns denotes p>0.05 (by unpaired Student’s t test or one-way ANOVA)

Given that ST2 is highly expressed in the uterine tissues of mice with endometrial injury, we hypothesized that IL33 might regulate the differentiation of ESCs into myofibroblasts via ST2 on ESCs. To validate this hypothesis, we used sST2 to block IL33 from binding to ST2 on ESCs during differentiation induction with TGF-β1. Our results demonstrated that the addition of sST2 reversed the inhibitory effects of activated LPMs on α-SMA protein expression (Fig. 5G) and downregulated the gene expression of *Acta2*, *Col1a1*, and *Kdm6b* (Fig. 5H). These findings indicated that activated LPMs inhibited the differentiation of ESCs into myofibroblasts by secreting IL33, which exerts its effects via the ST2-JMJD3 axis in ESCs.

### Result 6: Lars-Fos axis drives the IL33 expression of LPMs by binding to its enhancer

Given the elevated expression of IL33 in activated LPMs and its pivotal role in inhibiting inflammation and endometrial fibrosis, we sought to elucidate the molecular mechanisms driving IL33 expression. To this end, we conducted a functional enrichment analysis of DEGs using the Kyoto Encyclopedia of Genes and Genomes (KEGG) pathway (https://www.kegg.jp/). Our analysis revealed significant enrichment in pathways related to aminoacyl-tRNA ligase activity, cell adhesion molecule binding, and the aminoacyl-tRNA synthetase multienzyme complex (Fig. 6A). Gene Ontology (GO) analysis (https://geneontology.org/) indicated that DEGs upregulated at 6 hours post-AES treatment were mainly enriched in the Pentose Phosphate Pathway, HIF-1 signaling pathway, and Aminoacyl-tRNA biosynthesis (Fig. 6B). Gene Set Enrichment Analysis (GSEA) (https://www.gsea-msigdb.org/gsea/msigdb) also showed that upregulated genes were significantly associated with aminoacyl-tRNA ligase activity (Fig. 6C). Collectively, these results established a link between DEGs in the 6-hour group and aminoacyl-tRNA activity. A heatmap analysis of this pathway pinpointed leucyl-tRNA synthetase (Lars) as being notably upregulated at 6 hours (Fig. 6D). Since Lars mediates the activation of the mechanistic target of rapamycin complex 1 (mTORC1)^22^, we speculated that AES might regulate IL33 expression via Lars, as their expression patterns were closely aligned (Fig. 6E).

**Fig 6.**
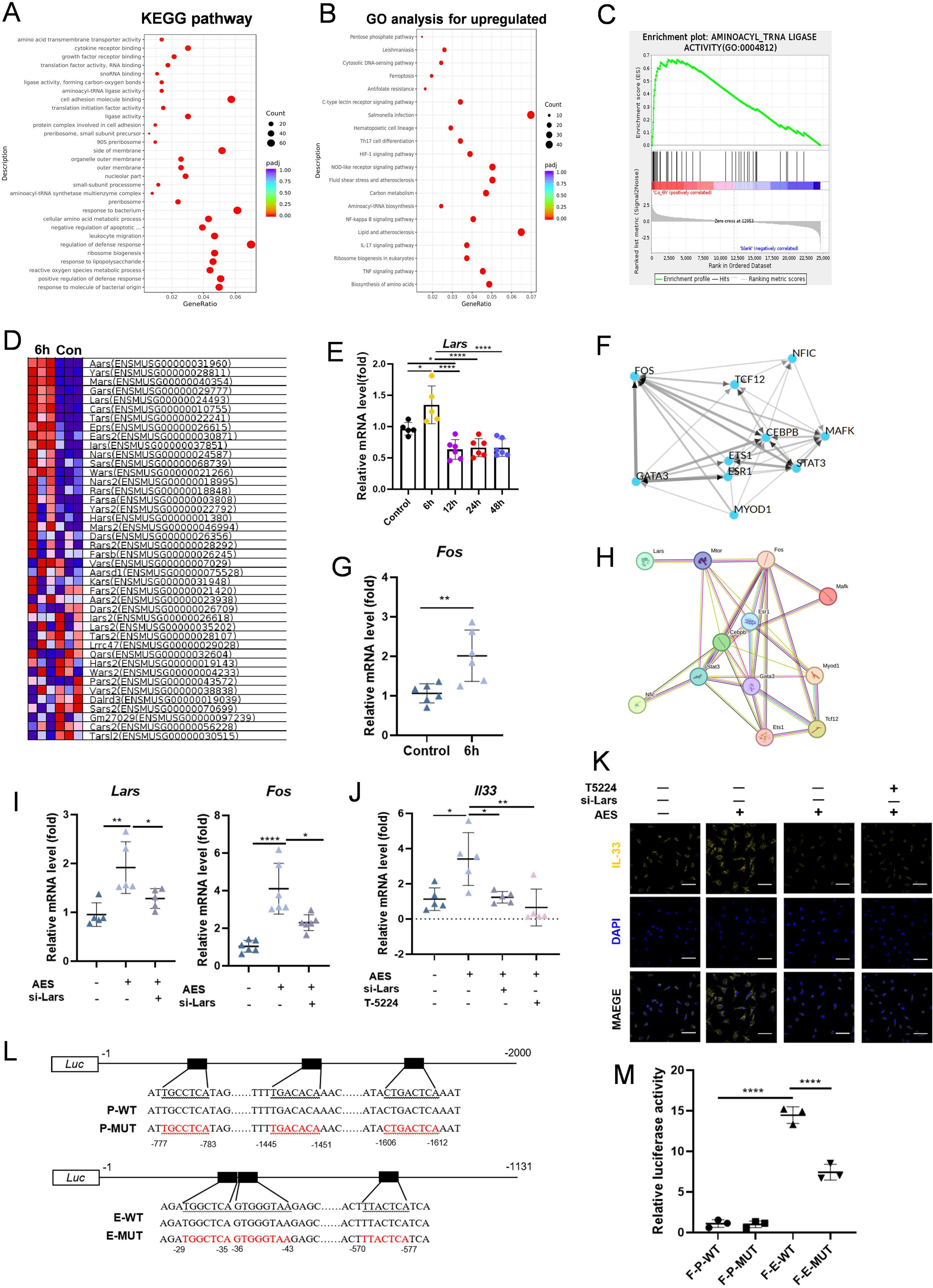
Lars-Fos axis upregulates Il33 expression in LPMs by modulating enhancers. A. KEGG enrichment analysis of all differentially up-expressed genes in 6h in LPMs after AES treated. B. GO enrichment analysis of all differentially up-expressed genes in 6h in LPMs after AES treated. C. GSEA enrichment analysis of all differentially up-expressed genes in 6h in LPMs after AES treated. D. The thermogram of AMINOACYL TRNA LIGASEACTIVITY pathway related genes expression pattern. E. The gene levels of *Lars* were detected with qRT-PCR (n=5-6). F. Interaction plot of TOP 10 transcription factors associated with up-regulation DEGs (ChEA3: https://maayanlab.cloud/chea3/). G. The gene levels of Fos were detected with qRT-PCR (n=6). H. Protein interactions between Lars and up-regulation DEGs-associated TOP10 transcription factors (String: https://cn.string-db.org/). I. The gene levels of *Lars* and *Fos* were detected with qRT-PCR (n=5-6). J. The gene levels of *Il33* were detected with qRT-PCR (n=5). K. The protein levels of Il33 were detected with immunofluorescence (n=3). (scale bar: 40 μm) L. Schematic diagram of plasmid construction. M. Dual luciferase reporter assay demonstrated that *Fos* positively regulates *Il33* expression by binding to enhancers (n=3). (F-P-WT: co-transfection of fos plasmid with promoter plasmid. F-P-MUT: co-transfection of fos plasmid with mutant promoter plasmid. F-E-WT: co-transfection of fos plasmid with enhencer plasmid. F-E-MUT: co-transfection of fos plasmid with mutant enhencer plasmid.) Values are mean±SD. *p<0.05, **p<0.01, ***p<0.001, ****p<0.0001, ns denotes p>0.05 (by unpaired Student’s t test or one-way ANOVA) N.

To explore the intricate molecular mechanisms, we utilized the ChEA3 database (https://maayanlab.cloud/chea3/) to identify the top 10 transcription factors (TFs) linked to upregulated DEGs in LPMs after 6 hours of AES treatment. Among these, Fos stood out as a pivotal TF, engaging in interactions with the remaining nine TFs (Fig. 6F). As the Fos family forms a tight complex with the Jun family to constitute the major component of AP-1, we integrated expression of Fos and Jun family members in our RNA-seq data and found the *Fos* was the significant expression gene in them (Supplementary Fig. 7A), a result further validated by qRT-PCR (Fig. 6G).

Additionally, by using the STRING database (https://cn.string-db.org/), we also identified a robust correlation between Fos and Lars (Fig. 6H), prompting us to postulate that Lars regulates Fos expression. Knockdown of *Lars* via siRNA resulted in a concomitant decrease in *Fos* gene expression (Fig. 6I). Moreover, either knocking down *Lars* or inhibiting Fos’s DNA-binding activity with T-5224 significantly reduced Il33 expression at both transcriptional and protein levels (Fig. 6J, K). These findings suggest that in LPMs, Lars modulates *Il33* gene expression by regulating Fos.

Fos typically promotes gene expression by binding to promoters or enhancers^23, 24^. Using the enhancer and promoter regions of *Il33* predicted by NCBI (https://www.ncbi.nlm.nih.gov/gene/), we conducted an analysis of these sequences within the UCSC Genome Browser (https://genome.ucsc.edu/) and identified potential binding sites for Fos. Further motif prediction using the JASPAR database (https://jaspar.elixir.no/) confirmed these findings (Supplementary Fig. 7B, C).

Dual-luciferase reporter assays demonstrated that the transfection of Fos significantly enhanced luciferase activity driven by the *Il33* enhancer region, and this effect was abolished when Fos-binding motifs were mutated (Fig. 6L-M). This phenomenon was not observed in the promoter of *Il33*. Collectively, these results indicated that AES upregulates Lars, which subsequently enhances Fos binding to the Il33 enhancer, thus promoting IL33 expression in LPMs.

## Discussion

In this study, we identified a unique subset of macrophages, termed GATA6^+^ macrophages or LPMs, that originally reside in the peritoneal cavity but subsequently migrate to the injured endometrium. Upon activation by injured endometrium, these LPMs exert an inhibitory effect on endometrial fibrosis by secreting IL33. Mechanistically, the upregulation of IL33 expression in activated LPMs is orchestrated by the Lars-Fos signaling axis, which binds to the enhancer region of the IL33 gene. IL33, produced by LPMs, suppresses the differentiation of ESCs into myofibroblasts by activating the ST2-JMJD3 signaling pathway in ESCs, thereby attenuating the fibrotic response in the endometrium.

GATA6^+^ macrophages, also referred to as LPMs in the peritoneal cavity, represent a subpopulation of macrophages that reside within the peritoneal, pleural, and pericardial cavities^3^. In their basal state, LPMs demonstrate swift mobility within the peritoneal cavity, partly attributed to diaphragmatic motion during respiration. However, in response to tissue or organ damage, LPMs are quickly recruited to the injury site, accumulate there within minutes^25^. For instance, following liver^26^ or gut^27^ injuries, LPMs migrate from the peritoneal cavity to the damaged region, initiating repair processes that promote revascularization and tissue regeneration. Similarly, GATA6^+^pericardial cavity macrophages (GPCMs) play a crucial role in cardiac repair and revascularization following myocardial infarction. Studies have shown that upon migration to the damaged heart, GPCMs experience a downregulation in GATA6 and CD102 expression without impairing their anti-fibrotic functions. However, the possibility of false-positive results in this context cannot be entirely excluded by the authors^16^. Our previous research demonstrated that mesenchymal stem cells (MSCs) facilitate the migration of LPMs to the uterus in a mouse model of IUA, thereby providing protective effects to the injured uterus^13^. However, the functional roles of LPMs following their migration to the damaged endometrium, as well as the precise mechanisms underlying these effects, remain largely unknown. In this study, through the analysis of clinical samples and murine experiments, we further characterized the migration of LPMs from the peritoneal cavity to the uterus during endometrial injury. Depletion of LPMs exacerbated the severity of the disease. By using fluorescent beads for the specific labeling of LPMs, we eliminated the possibility of false-positive results and observed phenotypic changes in LPMs after their migration to the damaged endometrial. RNA-seq revealed that activated LPMs displayed distinct transcriptional profiles, indicating a reprogramming of these cells upon activation, consistent with previous reports^16, 26^.

The role of IL33 in fibrosis continues to be an area of intense research. IL33 recombinant protein therapy at doses ranging from 0.4 µg to 1 µg, has demonstrated efficacy in alleviating fibrosis in multiple organs, including the heart^28^, kidneys^29^, and colon^30^, as well as promoting tissue repair in skin wounds^31, 32^. However, Liu et al. reported that intrauterine injection of a higher dose of IL33 (4 µg) exacerbated IUA^33^. This dose notably exceeds the therapeutic range for anti-fibrotic effects and the serum levels typically associated with disease pathology. In fact, doses within this higher range have been associated with the promotion of fibrosis ^34, 35^. These observations imply a dose-dependent effect of IL33 on fibrosis, with lower doses inhibiting fibrosis and promoting tissue repair, while higher doses may exacerbate it. In our study, intraperitoneal administration of IL33 at doses ranging from 0.25 µg to 1 µg effectively mitigated systemic inflammation and reduced endometrial fibrosis in a mouse model of endometrial injury. Furthermore, in clinical endometria with varying severities of IUA, we observed an upregulation of IL33 expression in GATA6^+^ macrophages, which diminished as disease severity increased. This suggests that IL33 levels secreted by LPMs are inversely correlated with the severity of IUA, further supporting the hypothesis that low doses of IL33 secreted by LPMs inhibit endometrial fibrosis.

However, our results do not suggest that simply replacing LPMs with low-dose recombinant IL33 injection represents the optimal therapeutic strategy for endometrial fibrosis. Although exogenous IL33 administration can be achieved through injections or drug delivery systems, LPMs may be essential for modulating the local IL33 microenvironment and fine-tuning its release. The direct application of IL33 poses challenges in precise dose regulation and may increase the risk of adverse effects, whereas IL33 secretion by LPMs could provide a more physiologically regulated approach to maintaining therapeutic levels. Furthermore, LPMs may play broader roles in tissue repair beyond IL33 secretion, potentially facilitating interactions with other immune cells and supporting additional regenerative processes^13, 36^. A compromised ability of LPMs to produce IL33, possibly due to underlying pathological conditions or immune dysfunction, could contribute to the pathogenesis in some patients. Investigating the molecular mechanisms governing IL33 secretion by LPMs could help identify patient subgroups who are most likely to benefit from IL33-based therapies or guide the development of personalized treatment regimens tailored to individual immune profiles.

The differentiation of ESCs into myofibroblasts is a critical step in endometrial fibrosis progression. Previous research has shown that inhibition of JMJD3 enhances TGFβ1 signaling in renal mesangial cells, thereby promoting their transition into myofibroblasts^21^. Our findings indicated that IL33 inhibits the differentiation of ESCs into myofibroblasts by binding to its receptor ST2, which subsequently upregulates JMJD3 expression in ESCs. The activation of the ST2-JMJD3 axis is essential for LPMs-derived IL33 to exert its inhibitory effect on endometrial fibrosis. Regarding the regulation of IL33 expression, we discovered that the Lars-FOS axis plays a pivotal role. Lars, known to activate the mTOR signaling pathway via mTORC1 activation^37^, was identified as a key regulator in this process. The mTOR pathway can modulate the expression of Fos^38^. For the first time, we reported that Lars upregulated IL33 expression in activated LPMs by regulating Fos, which binds to the enhancer region of the *Il33* gene.

The endometrium stands unique among mammalian tissues, possessing the remarkable capacity for complete functional regeneration after cyclical shedding. Under physiological conditions, the endometrium undergoes a series of proliferation, differentiation, and shedding phases, allowing for full functional repair without the formation of scar tissue^39^. Endometrial damage incurred during surgical procedures, such as curettage, shares similarities with natural endometrial shedding during menstruation^4, 40^. In both scenarios, the endometrial lining is shed, leading to cell death and the release of tissue fragments, which in turn release DAMPs, triggering a state of sterile inflammation^41^. In a mouse model mimicking menstruation, we found that LPMs migrated from the peritoneal cavity to the uterus during endometrial shedding induced by hormone withdrawal (Supplementary Fig. 8). Building upon this finding and our previous research, we propose that this migration of LPMs may contribute to the physiological, scar-free repair of the endometrium that is characteristic of menstruation in mammals.

In conclusion, our study revealed that DAMPs released by mechanically injured ESCs stimulate the migration of LPMs to the damaged uterus. These LPMs then upregulate IL33 expression via the Lars-Fos axis, achieved through binding to the enhancer region of the *Il33* gene. LPMs-derived IL33 inhibits the differentiation of ESCs into myofibroblasts by binding to ST2 receptor on the surface of ESCs. This interaction subsequently upregulates JMJD3 expression and suppresses fibrotic transformation, thereby preventing endometrial fibrosis and alleviating the severity of endometrial injury. Our findings underscore the pivotal role of LPMs in endometrial repair, with LPMs-derived IL33 playing a central role in maintaining endometrial homeostasis and preventing pathological fibrosis.

## Data availability

Source data are provided with this paper.

## Compliance with ethics requirements

This study was approved by the Ethics Committee of the Affiliated Drum Tower Hospital of Nanjing University (No. 2021-078-01) and all participants have provided informed consent before the endometrial biopsy. For animal experiments, all experiments involving animals were conducted according to the ethical policies and procedures approved by the Animal Protection and Ethics Committee of Nanjing University (IACUC-D2202077) followed the guidelines of the National Institutes of Health (NIH) for the care and use of laboratory animals. Every effort was made to minimize animal suffering and the number of animals used in the experiments. The authors declare that they have adhered to the ethical standards required for conducting research with human and animal subjects and have no conflicts of interest to disclose.

## Acknowledgements

We thank Yang Cheng (Cornell University, Department of Molecular Biology and Genetics) for her assistance and technical expertise. This work was supported by National Key R&D Program of China (2021YFC2701603), National Natural Science Foundation of China (82471663, 82271653, 82071600), Jiangsu Provincial Obstetrics and Gynecology Innovation Center (CXZX202229) and Jiangsu Biobank of Clinical Resources (BM2015004).

## Author contributions

L Yin: Conceptualization, Data curation, Investigation, Methodology, Investigation, Visualization, Writing - original draft

J Li: Methodology, Investigation, Visualization

Y Dong: Investigation, Visualization

J Wang, X Wang, Y Li, Y Hu: Investigation

Y Hou: Project administration, Supervision, Writing-review & editing

G Zhao: Conceptualization, Data curation, Funding acquisition, Project administration, Writing-review & editing

## Competing interests

The authors declare no competing interests.

**Correspondence and requests for materials** should be addressed to Guangfeng Zhao.

---

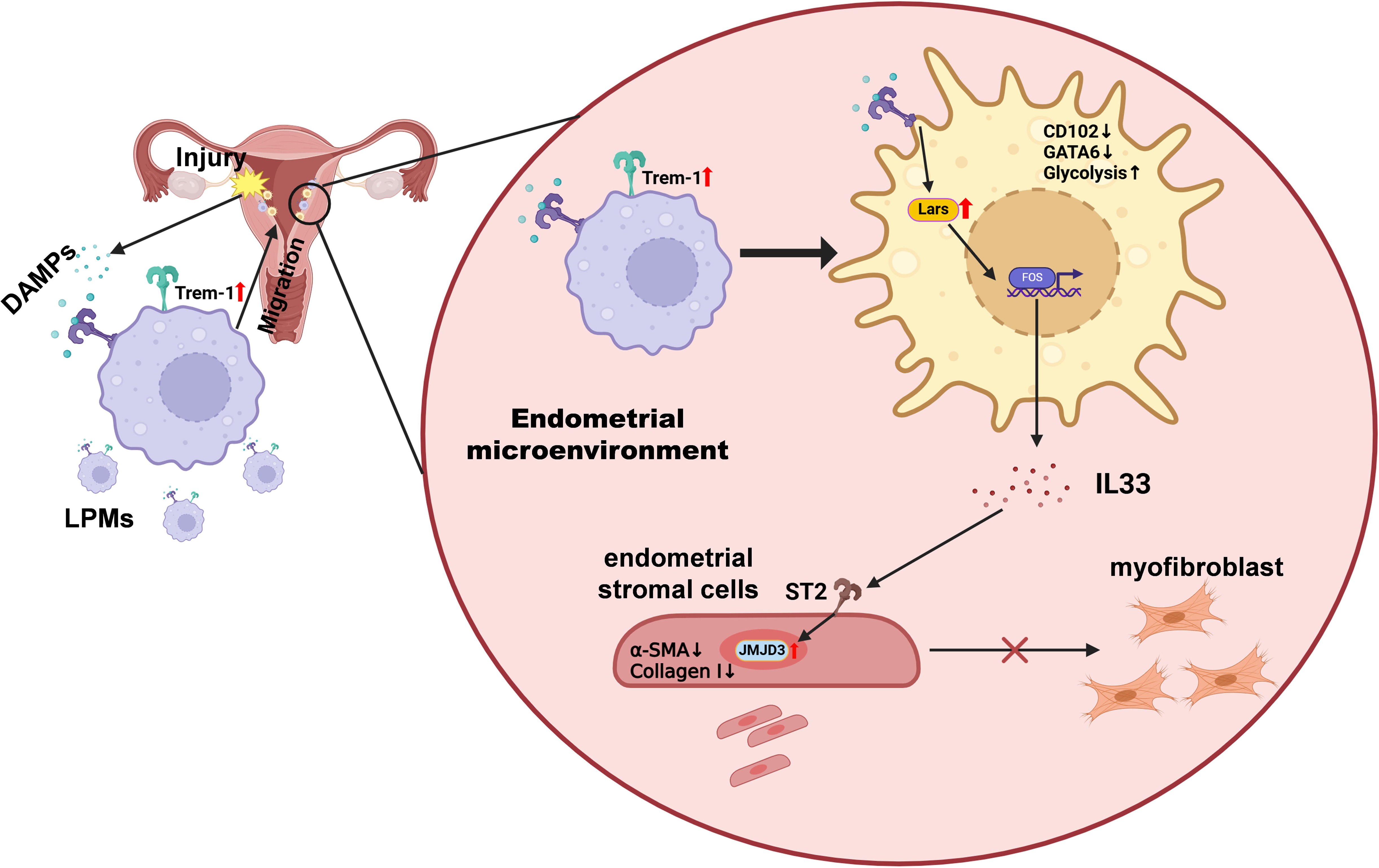

