## Supplementary figure & table for "Peritoneal cavity-derived GATA6^+^ macrophages inhibit fibrosis through IL33 in endometrium"

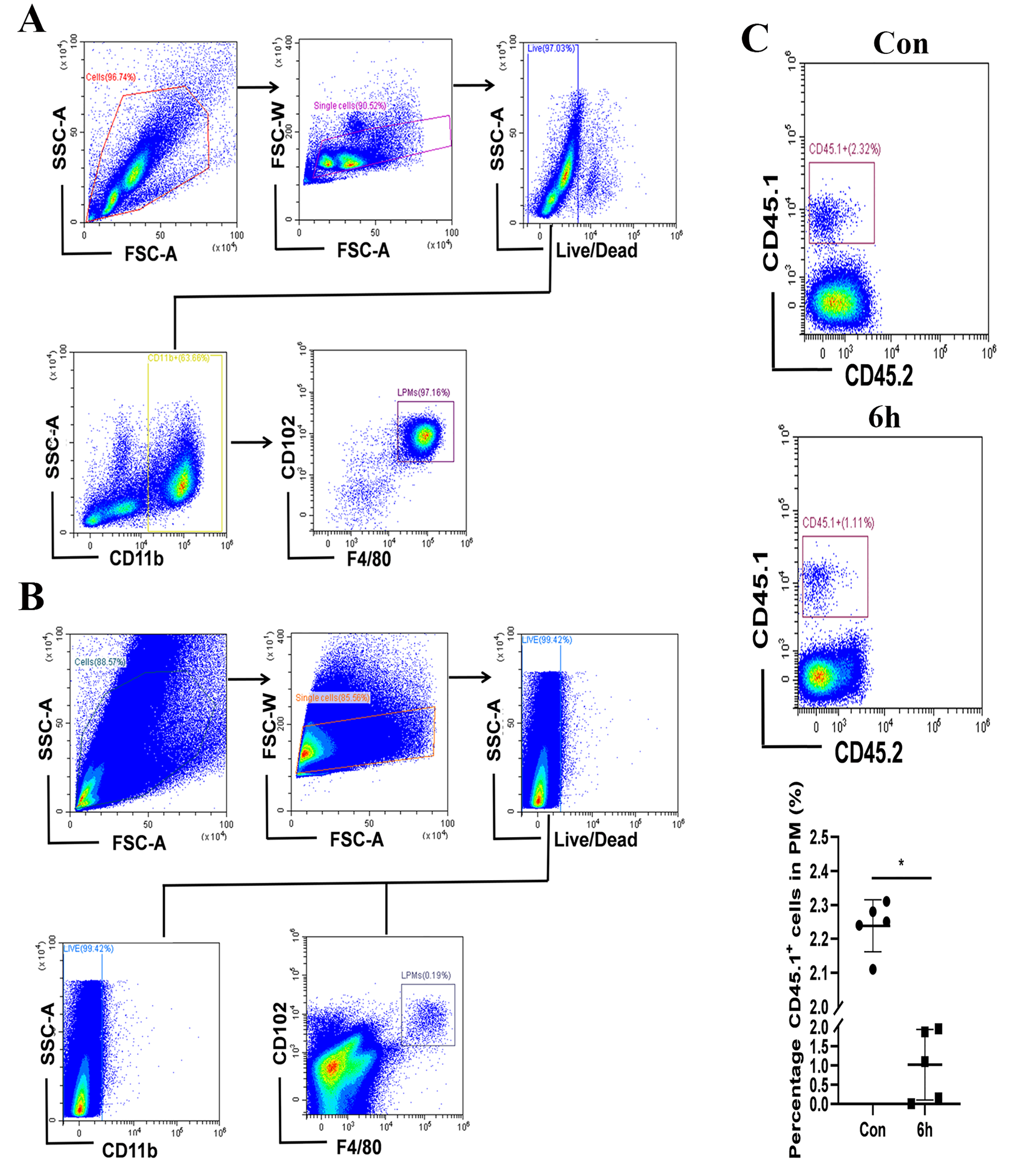

**Supplementary fig. 1**

A. The flow cytometry gating strategy of LPMs in the peritoneal cavity.

B. The flow cytometry gating strategy of LPMs in the uterus.

C. Flow cytometry analysis of changes in the proportion of CD45.1^+^ in the peritoneal cavity of normal control and endometrial injury mice (n=5)

Values are mean±SD. *p<0.05 (by unpaired Student’s t test)

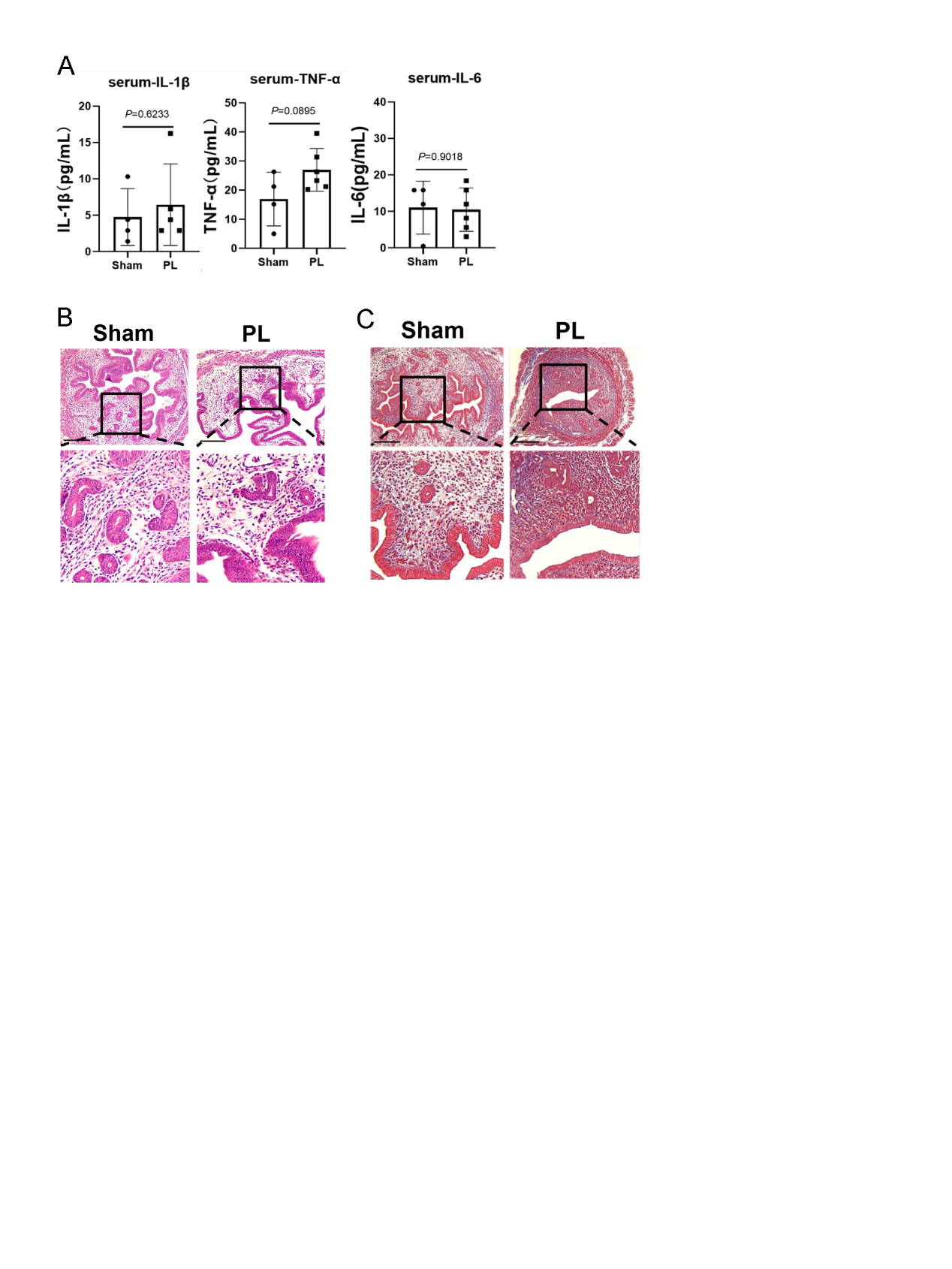

**Supplementary fig. 2**

A. Serum concentrations of IL-1β, TNF-α, and IL-6 were measured by ELISA (n=4-6).

B. HE staining of endometrial tissues obtained from mice (n=3).

C. Masson’s trichrome staining of endometrial tissues obtained from mice (n=3).

Scale bar indicates 200 μm. Values are mean±SD. ns denotes p>0.05 (by unpaired Student’s t test)

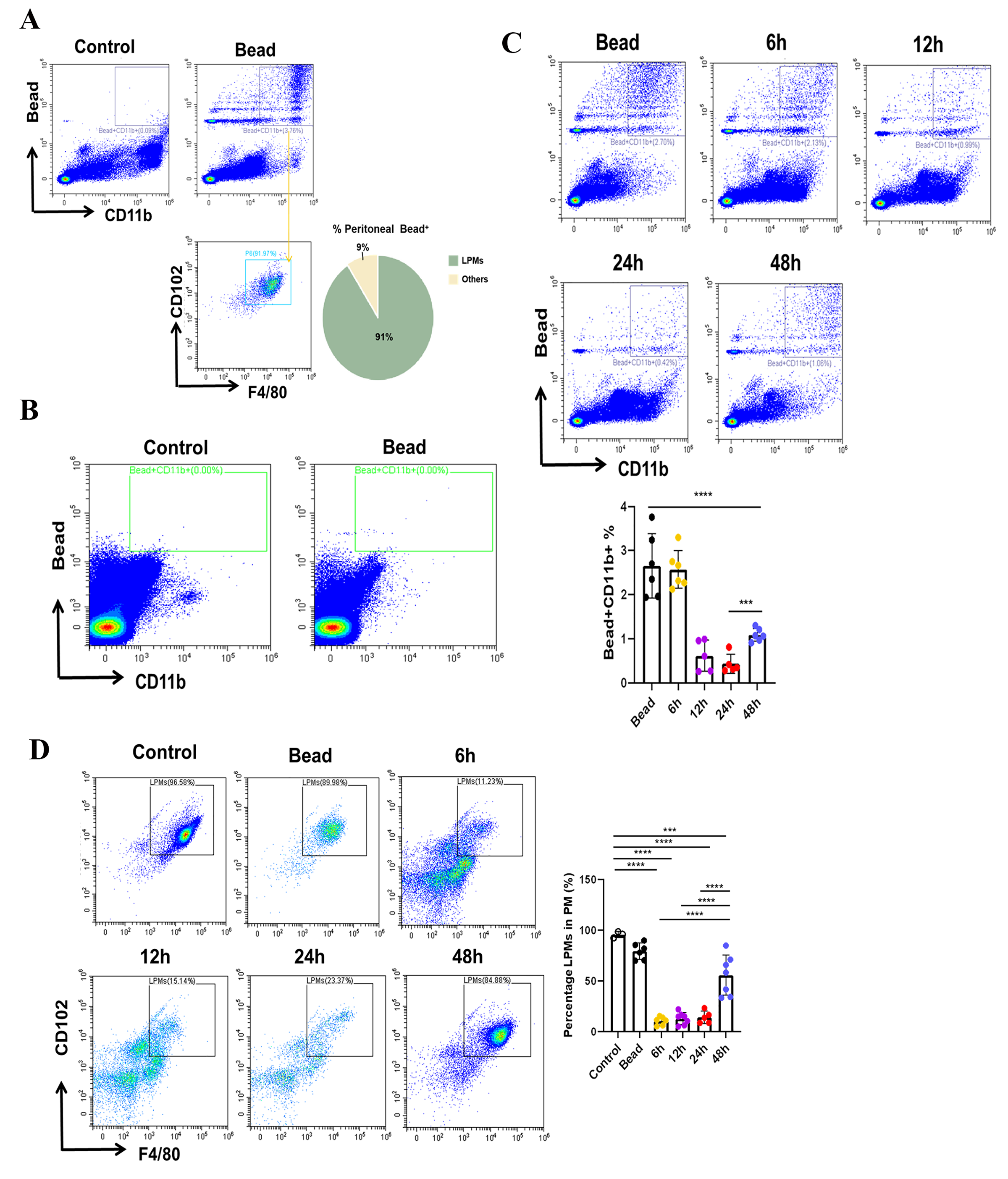

**Supplementary fig. 3**

A. The proportion of Bead specific labeled LPMs.

B. The effect of Bead on a healthy uterus.

C. Flow cytometry analysis of changes in the proportion of Beads^+^CD11b^+^cells and the change of them in the peritoneal cavity (n=5-6).

D. Flow cytometry analysis of changes in the proportion of LPMs and the change of them in the peritoneal cavity (n=5-6).

Values are mean±SD. *p<0.05, **p<0.01, ***p<0.001, ****p<0.0001, ns denotes p>0.05 (by unpaired Student’s t test or one-way ANOVA)

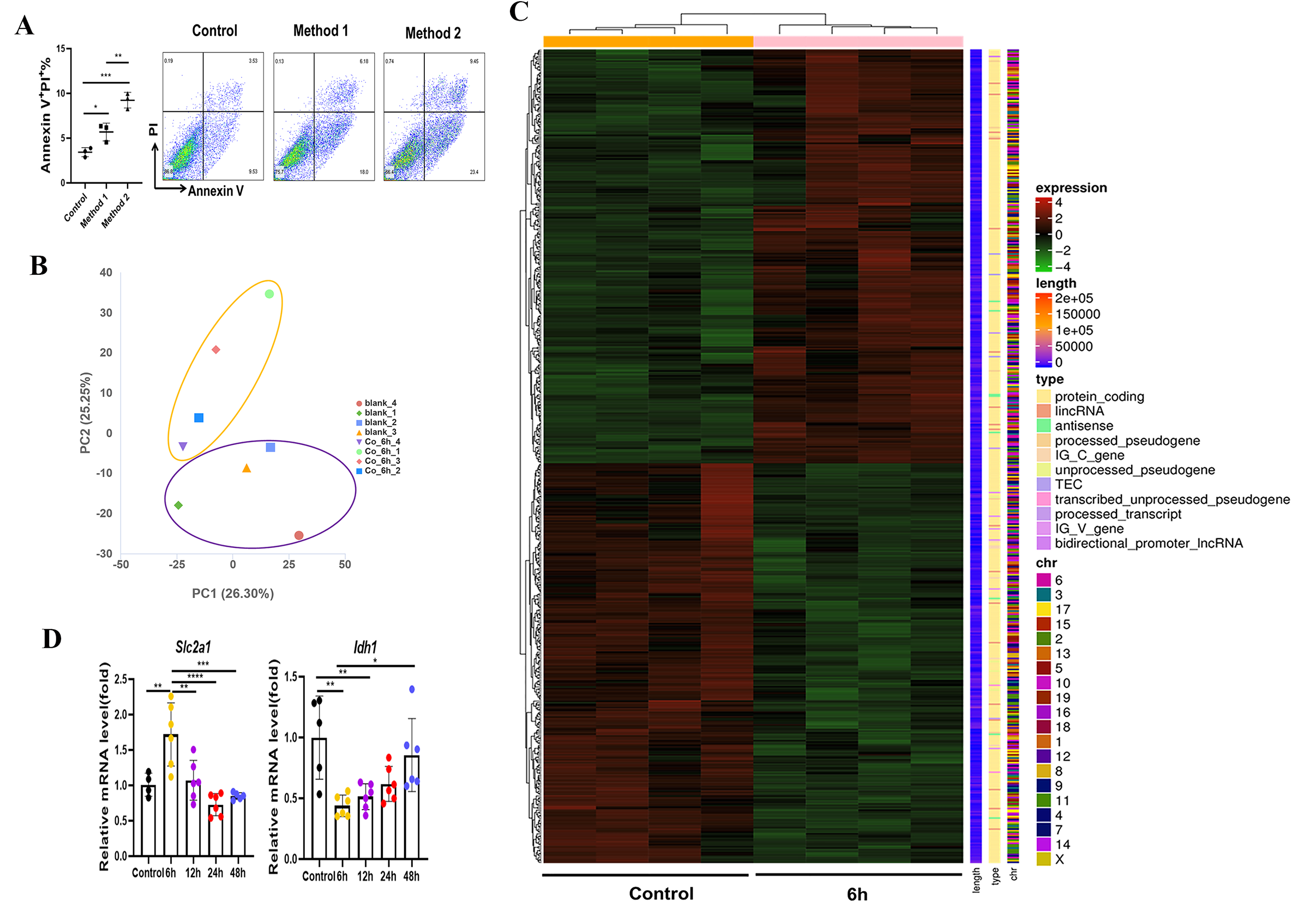

**Supplementary fig. 4**

A. Ratio of mechanical injury-induced apoptosis in ESCs detected by flow cytometry (n=3).

B. PCA analysis of samples.

C. The heatmap showing 541 up- and 646 down-regulated DEGs (p < 0.05 and absolute fold change >1) in endometria from AES-treated 6h (n = 4) and control (n = 4).

D. The gene levels of *Slc2a1* and *Idh1* were detected with qRT-PCR (n=5-6).

Values are mean±SD. *p<0.05, **p<0.01, ***p<0.001, ****p<0.0001, ns denotes p>0.05 (by unpaired Student’s t test or one-way ANOVA)

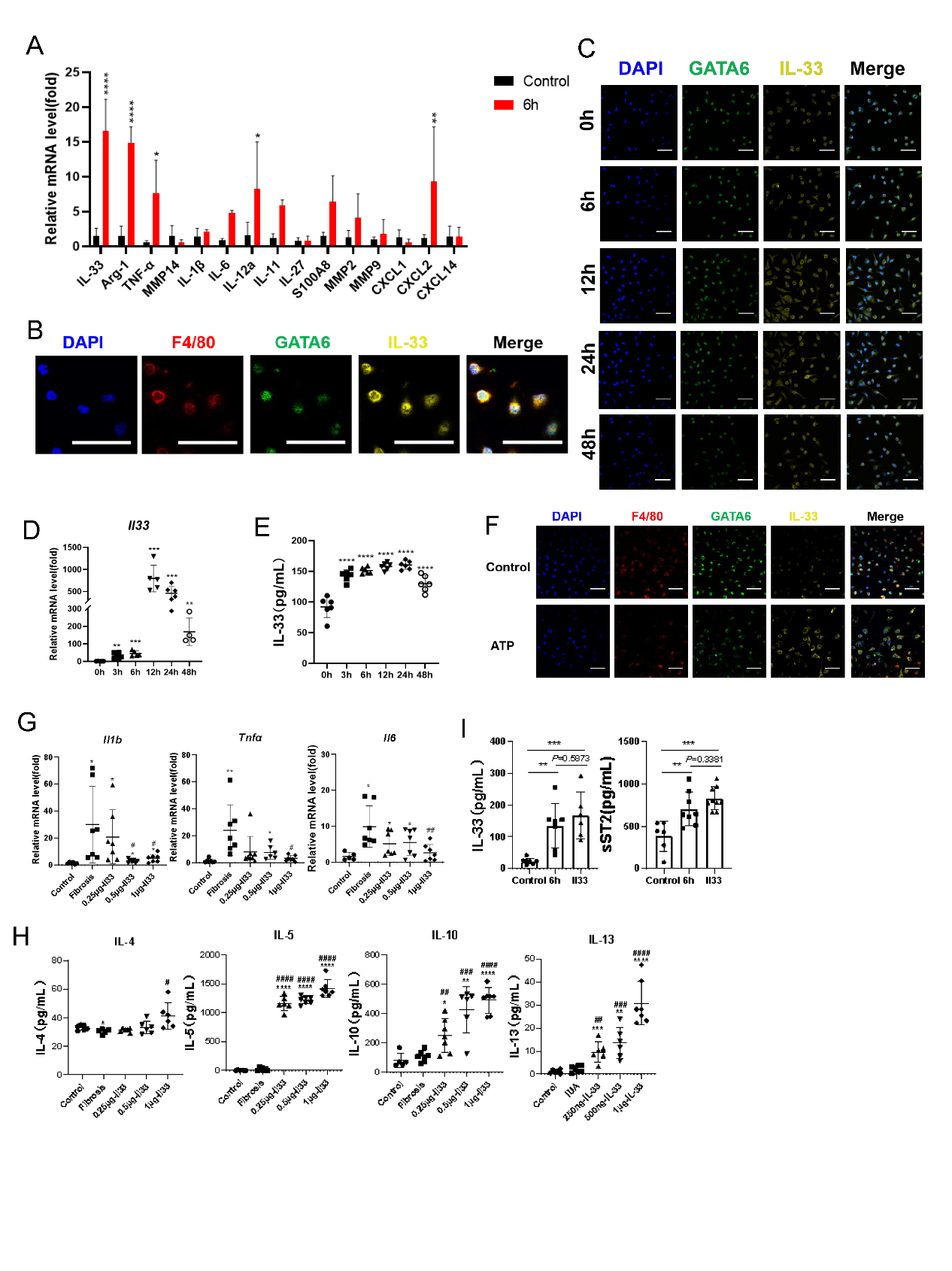

**Supplementary fig. 5**

A. The gene levels of differentially expressed cytokines were detected with qRT-PCR (n=3-4).

B. Representative immunostaining images of F4/80, GATA6 and IL33 from samples of LPMs (scale bar: 40 μm).

C. Representative immunostaining images of GATA6 and IL33 from samples of LPMs (scale bar: 40 μm).

D. The gene levels of *Il33* were detected with qRT-PCR (n=4-6)

E. Serum concentrations of Il33 were measured by ELISA (n=6)

F. Representative immunofluorescence images of F4/80, GATA6 and IL33 from samples of LPMs (scale bar: 40 μm).

G. The gene levels of *Tnfα*, *Il1b*, and *Il6* were detected with qRT-PCR (n=7).

H. Serum concentrations of IL-4, IL-5, IL-10, and IL-13 were measured by ELISA (n=6).

I. Serum concentrations of IL-33, and sST2 were measured by ELISA (n=6-7).

Values are mean±SD. *p<0.05, **p<0.01, ***p<0.001, ****p<0.0001, ns denotes p>0.05 (by unpaired Student’s t test or one-way ANOVA)

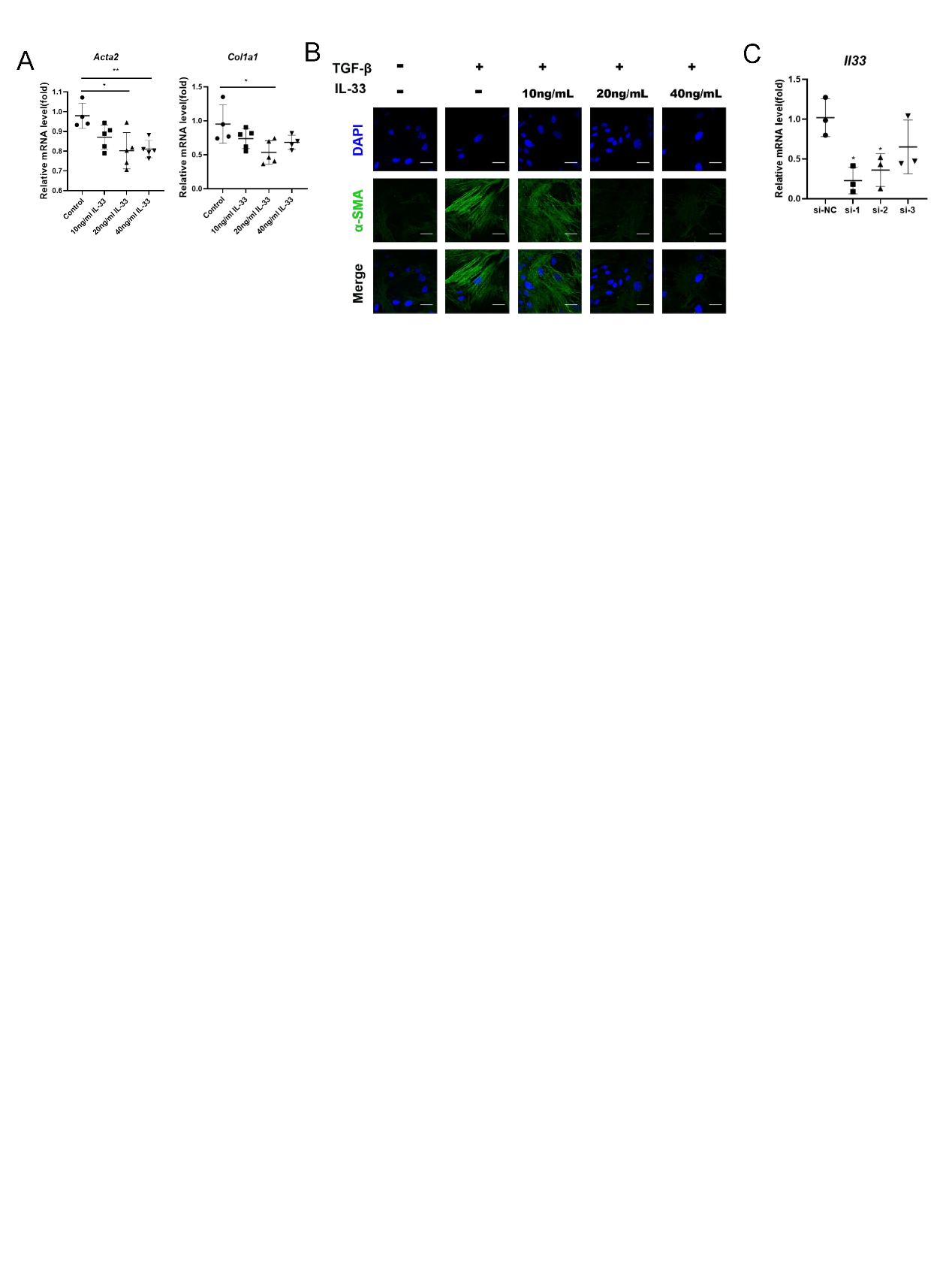

**Supplementary fig. 6**

A. The gene levels of *Acta2* and *Col1a1* were detected with qRT-PCR (n=4-5).

B. Representative immunofluorescence images of α-SMA from samples of ESCs (n=3) (scale bar: 100 μm).

C. The gene levels of *Il33* were detected with qRT-PCR (n=3).

Values are mean±SD. *p<0.05, **p<0.01, ***p<0.001, ****p<0.0001, ns denotes p>0.05 (by unpaired Student’s t test or one-way ANOVA)

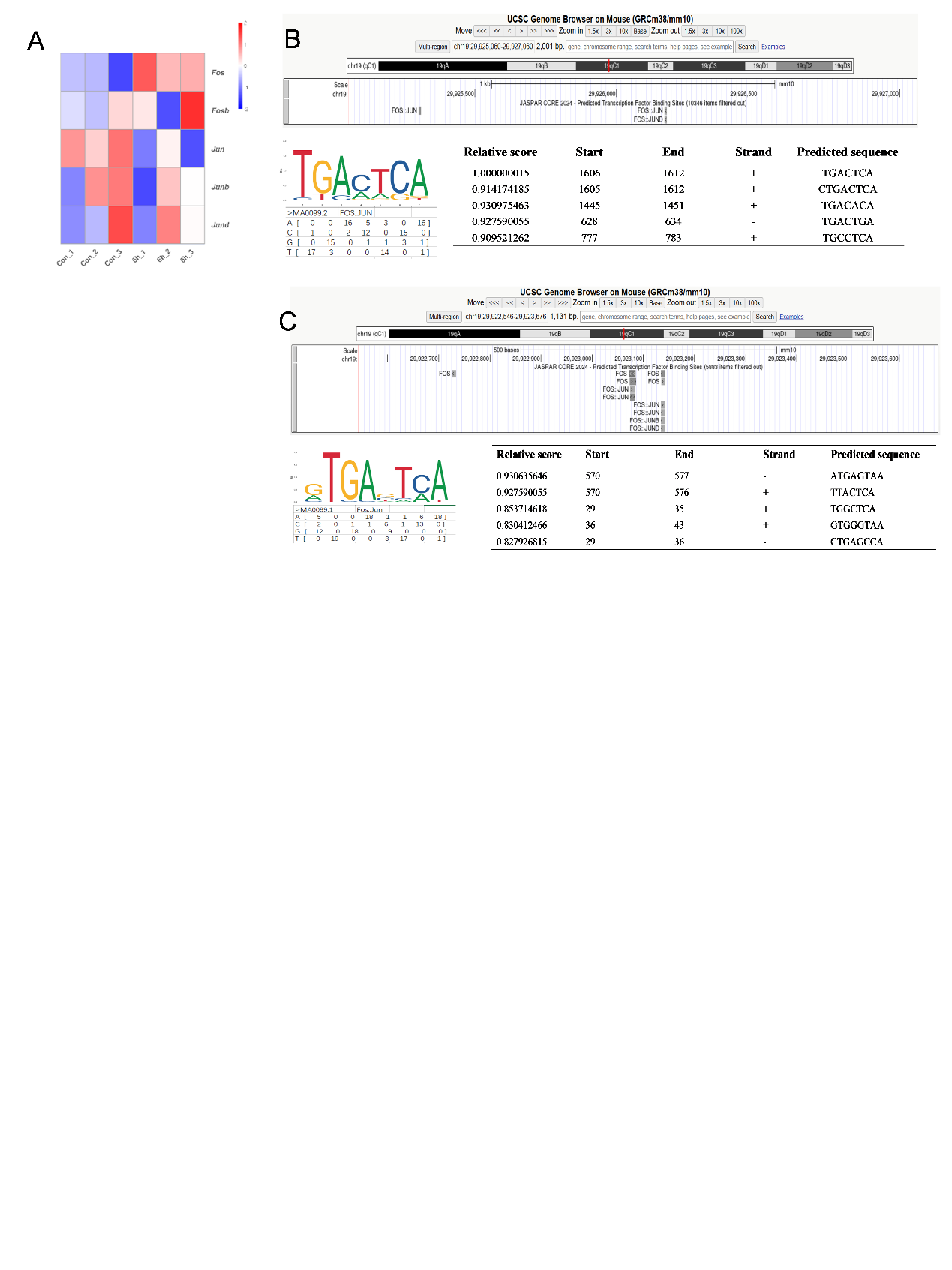

**Supplementary fig.7**

A. The thermogram of AP1 family related genes expression pattern.

B. Selection of Fos binding motifs on the *Il33* promoter region.

C. Selection of Fos binding motifs on the *Il33* enhencer region.

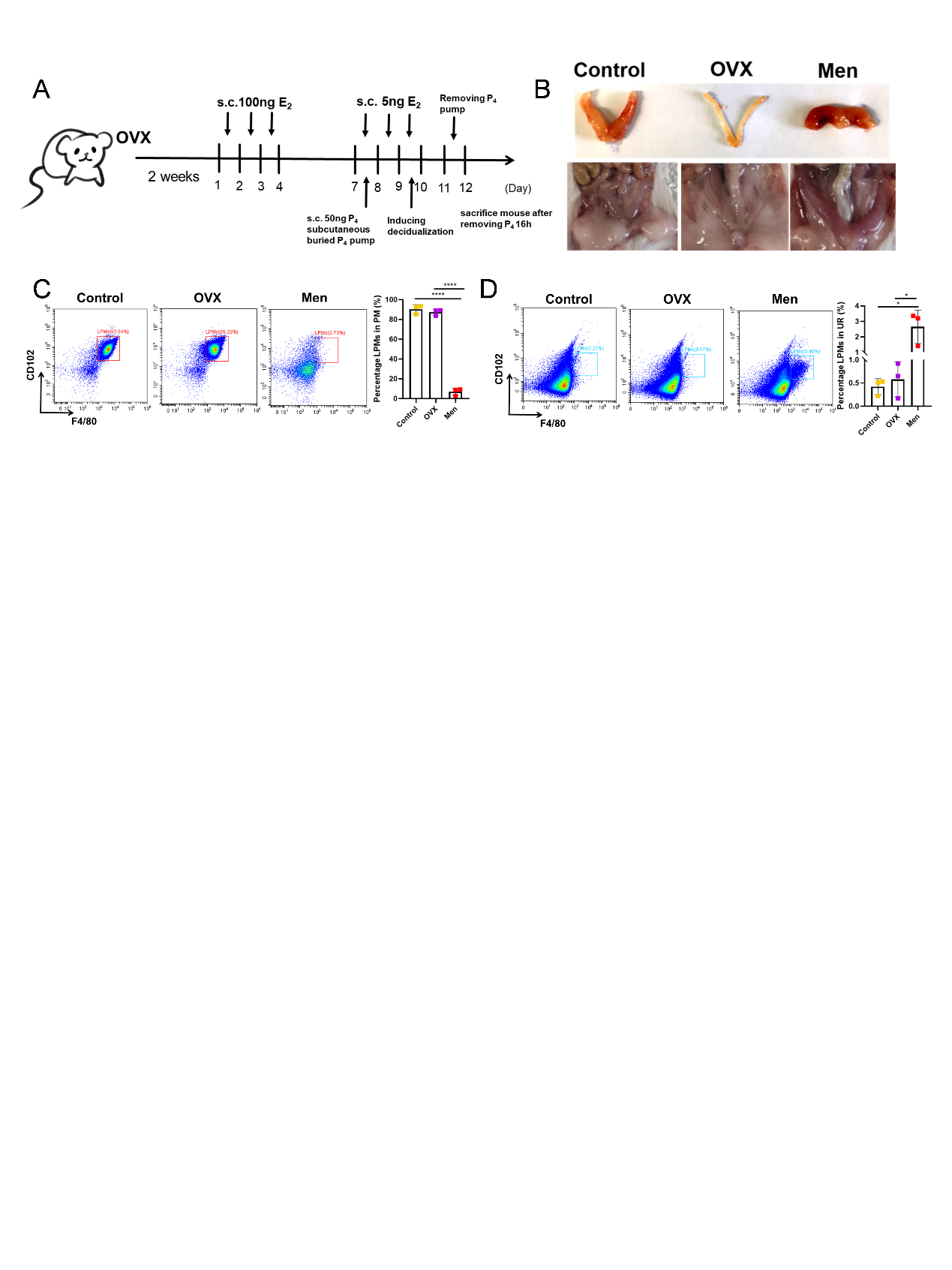

**Supplementary fig. 8**

A. Schematic diagram of the mouse menstrual model modeling.

B. Photograph of the uterus of a menstruating or non-menstruating mouse.

C. Flow cytometry analysis of changes in the proportion of LPMs in the mice peritoneal cavity (n=3). (OVX: ovariectomy mouse. Men: menstruating mouse)

D. Flow cytometry analysis of changes in the proportion of LPMs in the mice uterus (n=3).

Values are mean±SD. *p<0.05, **p<0.01, ***p<0.001, ****p<0.0001, ns denotes p>0.05 (by unpaired Student’s t test or one-way ANOVA)

### Supplementary Tables

**Table S1.** **Sequences of forward and reverse primers used for PCR amplification**

| **Gene** | **Forward Primer** | **Reverse Primer** |
| --- | --- | --- |
| *β-actin* | AGGTGACAGCATTGCTTCTG | GGGAGACCAAAGCCTTCATA |
| *Il1b* | TTCAGGCAGGCAGTATCACTC | GAAGGTCCACGGGAAAGACAC |
| *Il6* | TAGTCCTTCCTACCCCAATTTCC | TTGGTCCTTAGCCACTCCTTC |
| *Tnf* | CCACCACGCTCTTCTGTCTA | GGTCTGGGCCATAGAACTGA |
| *Gata6* | TTGCTCCGGTAACAGCAGTG | GTGGTCGCTTGTGTAGAAGGA |
| *Icam2* | TGGTCCGAGAAGCAGATAGTAG | GAGGCTGGTACACCCTGATG |
| *Trem1* | GACTGCTGTGCGTGTTCTTTG | GCCAAGCCTTCTGGCTGTT |
| *Il33* | TCCAACTCCAAGATTTCCCCG | CATGCAGTAGACATGGCAGAA |
| *Lars* | GAGCAGCAAGGGCAAATACTT | ACTGCAAACTCACACTTGGATAA |
| *Fos* | CGGGTTTCAACGCCGACTA | TTGGCACTAGAGACGGACAGA |
| *Ki67* | ATCATTGACCGCTCCTTTAGGT | GCTCGCCTTGATGGTTCCT |
| *Acta2* | GCTCCTCTTAGGGGCCACT | CCACGTCTCACCATTGGGG |
| *Col1a1* | TAAGGGTCCCCAATGGTGAGA | GGGTCCCTCGACTCCTACAT |
| *Kdm6b* | TGAAGAACGTCAAGTCCATTGTG | TCCCGCTGTACCTGACAGT |
| *Il27* | CTGTTGCTGCTACCCTTGCTT | CACTCCTGGCAATCGAGATTC |
| *Il11* | TGTTCTCCTAACCCGATCCCT | CAGGAAGCTGCAAAGATCCCA |
| *Mmp14* | CAGTATGGCTACCTACCTCCAG | GCCTTGCCTGTCACTTGTAAA |
| *Mmp9* | CTGGACAGCCAGACACTAAAG | CTCGCGGCAAGTCTTCAGAG |
| *Mmp2* | CAAGTTCCCCGGCGATGTC | TTCTGGTCAAGGTCACCTGTC |
| *Cxcl1* | CTGGGATTCACCTCAAGAACATC | CAGGGTCAAGGCAAGCCTC |
| *Cxcl2* | CCAACCACCAGGCTACAGG | GCGTCACACTCAAGCTCTG |
| *Cxcl14* | GAAGATGGTTATCGTCACCACC | CGTTCCAGGCATTGTACCACT |
| *S100a8* | AAATCACCATGCCCTCTACAAG | CCCACTTTTATCACCATCGCAA |
| *Il12a* | CTGTGCCTTGGTAGCATCTATG | GCAGAGTCTCGCCATTATGATTC |
| *Arg1* | CTCCAAGCCAAAGTCCTTAGAG | AGGAGCTGTCATTAGGGACATC |
| *Slc2a1* | CAGTTCGGCTATAACACTGGTG | GCCCCCGACAGAGAAGATG |
| *Idh1* | ATGCAAGGAGATGAAATGACACG | GCATCACGATTCTCTATGCCTAA |

**Table S2. Reagents and Sources**

| **REAGENT** | **SOURCE** | **IDENTIFIER** |
| --- | --- | --- |
| **Enzyme-linked immunosorbent assay Kits** | | |
| IL-1β | Biolegend | 432604 |
| IL-4 | Biolegend | 431104 |
| IL-5 | Biolegend | 431201 |
| IL-6 | Biolegend | 431304 |
| IL-10 | Biolegend | 431414 |
| IL-13 | Invitrogen | 335585-003 |
| IL-33 | Invitrogen | 349644-009 |
| TNFα | Biolegend | 430901 |
| sST2 | Multisciences | EK2163-96 |
| **Antibodies** | | |
| FITC anti-mouse IL-33 | R&D | IC3626J |
| PE anti-mouse L-33 | R&D | IC3626P |
| BB700 anti-mouse CD102 | Biosciences | 742107 |
| PE anti-mouse CD102 | Invitrogen | A15451 |
| APC anti-mouse F4/80 | Invitrogen | 17-4801-82 |
| PE-cy7 anti-mouse CD11b | Biolegend | 101216 |
| PE anti-mouse CD45.1 | Biolegend | 110707 |
| APC anti-mouse CD45.2 | Biolegend | 109813 |
| PE anti-mouse Ki67 | Biolegend | 652425 |
| PE anti-mouse GATA6 | Cell Signaling Technology | 26452S |
| IL33 | Proteintech | 66235-1-Ig |
| GATA6 | Proteintech | 55435-1-AP |
| GATA6 | R&D | AF1700 |
| CD102(ICAM2) | Abcam | ab189463 |
| ST2 | Proteintech | 11920-1-AP |
| α-SMA | Boster Biological Techonlogy | BM0002 |
| Vimentin | Proteintech | 10366-1-AP |
| CD68 | Cell Signaling Technology | 97778 |
| JMJD3 | Proteintech | 55334-1-AP |
| CD31 | Cell Signaling Technology | 77699S |
| F4/80 | Cell Signaling Technology | 30325T |
| Ki67 | Proteintech | 28074-1-AP |
| Anti-rabbit IgG, HRP-linked Antibody | Cell Signaling Technology | 7074S |
| Anti-mouse IgG, HRP-linked Antibody | Cell Signaling Technology | 7076S |
| GAPDH | Proteintech | 60004-1-Ig |
| **Chemicals, inhibitors, and Recombinant Proteins** | | |
| IL33 | Biolegend | 580514 |
| TGFβ1 | MCE | HY-P78360 |
| sST2 | MCE | HY-P72459 |
| T5224 | MCE | HY-12270 |
| BAY876 | MCE | HY-100017 |
| Estradiol | MCE | HY-B0141 |
| Progesterone | VETEC | V900699-5G |
| Clophosome-A and Control Liposomes | FormuMax | F70101C-AC-2 |
| Fluoresbrite YG Microspheres | PolysciencesInc. | 17154-10 |
| Lipofectamine™ 3000 Transfection Reagent | Invitrogen | [L3000015](https://www.thermofisher.cn/order/catalog/product/cn/en/L3000015) |

**Table S3.** **Sequences of siRNA**

| **siRNA name** | **Sequences (5’-3’)** |
| --- | --- |
| si-*Il33* | GGTGCTACTACGCTACTAT |
| si-*Lars* | CAGGCUGCGAAAGACAAAUTT |
| si-*Trem1* | GAGCGUCCCAUCCUUAUUATT |
